# Treatment with IFB-088 improves neuropathy in CMT1A and CMT1B mice

**DOI:** 10.1101/2021.10.18.464779

**Authors:** Yunhong Bai, Caroline Treins, Vera G. Volpi, Cristina Scapin, Cinzia Ferri, Rosa Mastrangelo, Thierry Touvier, Francesca Florio, Francesca Bianchi, Ubaldo Del Carro, Frank F. Baas, David Wang, Pierre Miniou, Philippe Guedat, Michael E. Shy, Maurizio D’Antonio

## Abstract

Charcot Marie Tooth diseases type 1A (CMT1A), caused by duplication of Peripheral Myelin Protein 22 (*PMP22*) gene, and CMT1B, caused by mutations in myelin protein zero (*MPZ*) gene are the two most common forms of demyelinating CMT (CMT1) and no treatments are available for either. Prior studies of the *Mpz*Ser63del mouse model of CMT1B have demonstrated that protein misfolding, endoplasmic reticulum (ER) retention and activation of the unfolded protein response (UPR) contributed to the neuropathy. Heterozygous patients with an arginine to cysteine mutation in MPZ (*MPZ*R98C) develop a severe infantile form of CMT1B which is modeled by *Mpz*R98C/+ mice that also show ER-stress and an activated UPR. C3-PMP22 mice are considered to effectively model CMT1A. Altered proteostasis, ER-stress and activation of the UPR have been demonstrated in mice carrying *Pmp22* mutations. To determine whether enabling the ER-stress/UPR and readjusting protein homeostasis would effectively treat these models of CMT1B and CMT1A we administered Sephin1/IFB-088/icerguestat, a UPR modulator which showed efficacy in the MpzS63del model of CMT1B, to heterozygous *Mpz*R98C and C3-PMP22 mice. Mice were analyzed by behavioral, neurophysiological, morphological and biochemical measures. Both *Mpz*R98C/+ and C3-PMP22 mice improved in motor function and neurophysiology. Myelination, as demonstrated by g-ratios and myelin thickness, improved in CMT1B and CMT1A mice and markers of UPR activation returned towards wild type values. Taken together our results demonstrate the capability of IFB-088 to treat a second mouse model of CMT1B and a mouse model of CMT1A, the most common form of CMT. Given the recent benefits of IFB-088 treatment in Amyotrophic Lateral Sclerosis and Multiple Sclerosis animal models, these data demonstrate its potential in managing UPR and ER-stress for multiple mutations in CMT1 as well as in other neurodegenerative diseases.

## Introduction

Charcot Marie Tooth (CMT) disease refers to heritable peripheral neuropathies, which are the most common genetic neuromuscular diseases, affecting 1:2500 individuals [60]. Autosomal dominant (AD) inheritance is the most common, followed by X-linked and autosomal recessive (AR) forms. Most forms of CMT are demyelinating while approximately one-third appear to be primary axonal disorders [18, 43]. CMT1A is the most common form, affecting approximately half of all patients with CMT [18], and is caused by a 1.4Mb duplication within chromosome 17p11.2, in the region containing the peripheral myelin protein 22 (*PMP22)* gene [39, 63]. CMT1B (caused by mutations in myelin protein zero (*MPZ)* gene, encoding for P0 protein) [26] is the second most frequent AD demyelinating form, encompassing around 5% of CMT cases [18]. At present there are no effective treatments to slow progression or improve neuropathy in patients with CMT1A or CMT1B.

80% of newly synthesized PMP22 is rapidly degraded by the proteasome under normal conditions, with only 20% reaching the cell surface or myelin sheath [48]. Whether this ratio is altered when PMP22 is over-expressed in CMT1A is unknown. Ordinarily, the maintenance of correct protein homeostasis is tightly controlled by protein quality control mechanisms [1]. The elevated protein expression associated with *PMP22* trisomy, in conjunction with the instability of the PMP22 protein, impose a heavy burden on the endoplasmic reticulum (ER) protein quality control (ERQC) mechanisms [17, 35, 45, 57]. When these control mechanisms fail, stress responses are activated leading to protein-kinase RNA-like endoplasmic reticulum kinase (PERK) (one of the unfolded protein response (UPR) sensors) mediated phosphorylation of the alpha subunit of the eukaryotic translation initiation factor 2 (eIF2α)[25]. This phosphorylation causes an attenuation of global protein synthesis which reduces protein overload in the ER while allowing the translation of selected genes supporting stress recovery [38]. The extent to which these responses are activated in CMT1A is unclear, though there have been studies suggesting UPR activation in a CMT1A mouse model carrying seven copies of the human *PMP22* gene, the C22 mouse, which shows a severe dysmyelinating neuropathy [23], and in Trembler J (Tr^J^) mice [46], which are caused by a missense mutation in *Pmp22* [33].

Over 200 different mutations in *MPZ* cause neuropathy (http://hihg.med.miami.edu/code/http/cmt/public_html/index.html#/) and the disease mechanisms are largely unknown. One group of mutations presents clinically in infancy or early childhood, with very slow nerve conduction velocities (NCV), and dysmyelination morphologically. Another group of *MPZ* mutations does not present clinically until adulthood, with near normal NCV, and axonal damage but minimal demyelination morphologically (sometimes called CMT2I or CMT2J) [55]. Several *MPZ* mutations in these groups have been shown *in vitro* to cause the accumulation of the mutant protein within the ER, where it triggered ER stress and activated the UPR. These included *Mpz* S51ΔW57 [21], 506delT and 550del3insG [4, 32]. Moreover, two mouse models of CMT1B, the *Mpz*Ser63del and *Mpz*R98C mice [53, 56], demonstrated ER retention of the mutant protein and a canonical UPR. The compound IFB-088 (icerguastat, also known as sephin1) was developed to prolong protein translation attenuation in response to stress by inhibiting eIF2α dephosphorylation to allow cells to restore cellular homeostasis [12]. Oral treatment with IFB-088 largely prevented the molecular, motor and morphological abnormalities of the neuropathy of *Mpz*Ser63del mice [12]. Whether treatment with IFB-088 is effective for only the Ser63del *MPZ* mutation is not known. This is an important question perhaps since as many as 40% of *MPZ* mutations have been recently shown to activate components of the UPR [4]. Accordingly, we elected to treat *Mpz*R98C/+ mice with IFB-088.

We also hypothesized that IFB-088 may have effects on CMT1A, because of the highly metastable nature of PMP22 and because of UPR activity in C22 and Tr^J^ mice [23, 33]. We therefore chose to use IFB-088 to treat the C3-PMP22 mouse model of CMT1A. C3-PMP22 mice, which carry 3-4 copies of the human *PMP22* gene, develop a slowly progressive dysmyelinating peripheral neuropathy that is thought to be an appropriate model of CMT1A [15, 65]. Both CMT1B and CMT1A models were treated IFB-088, and evaluated by behavioral, neurophysiological, morphological and biochemical analysis.

## Materials and methods

### Myelinating DRG explant cultures

Dorsal-root-ganglia (DRG) were dissected from embryos at embryonic day 13.5 (E13.5) and plated singularly on collagen-coated coverslips as previously described [62]. Myelination was induced with 50 µg/ml ascorbic acid (Sigma Aldrich). Treatment with IFB-088 at the indicated concentration was applied for 2-week in parallel to the induction of myelination. Samples were then fixed and rat anti-MBP (1/5) [11] and rabbit anti-NF-H (1/1000, EMD Millipore) primary antibodies were added o/n at 4°C. The following day, DRGs were washed and FITC- or TRITC-conjugated secondary antibodies (1:200, Cappel) were added for 1h at room temperature. Specimen were incubated with DAPI (1:1000, SIGMA) and mounted with VectaShield (Vector Laboratories). 8-10 images were taken from each DRG using a fluorescence microscope (Leica DM5000) with a 10x objective, and the number of MBP+ internodes in each image was counted.

### Animal models

All experiments involving animals were performed in accord with experimental protocols approved by the San Raffaele Scientific Institute and the University of Iowa Animal Care and Use Committee. *Mpz*S63del transgenic mice [70] and *Mpz*R98C/+ knock-in mice [56] were maintained on the FVB/N background. C3-PMP22 transgenic mice [65] were obtained from the Amsterdam University Medical Center, Amsterdam, the Netherland. They were maintained on a C57BL/6J background. C3-PMP22 cohorts for this study were generated via *in vitro* fertilization following the protocol from the European Mouse Mutant Archive (EMMA) - mouse sperm cryopreservation protocol [61].

### Experimental design

#### C3-PMP22 study

Starting from post-natal day (PND) 15, mice were administered via oral gavage twice a day (bis in die (*b*.*i*.*d*.)). WT mice were administered with vehicle (saline solution: NaCl 0.9%) *b*.*i*.*d*. and C3-PMP22 mice were administered with vehicle *b*.*i*.*d*. or IFB-088 0.5mg/kg *b*.*i*.*d*. or 1mg/kg *b*.*i*.*d*. After 10-week treatment period, mice were tested for treadmill and grip-strength. After 12-week treatment period, mice were analysed for neurophysiology and sacrificed for morphology, and biochemistry. Each experiment was performed by a different operator, completely blinded to genotype and treatment.

#### *Mpz*R98C/+ study

WT and *Mpz*R98C/+ mice were administrated via oral gavage twice a day with vehicle (saline solution: NaCl 0.9%) or IFB-088 1mg/kg starting from PND30. After 90 days (PND120) and 150 days (PND180) of treatment, mice were tested for rotarod, grip-strength and electrophysiology. Animals were then sacrificed for morphology, protein, and molecular expression analysis with the evaluator blinded to genotype and treatment.

### Treadmill

C3-PMP22 mice were evaluated on treadmills after 10-week treatment. A grid that delivers a mild electric shock is used to motivate the animal to run. On the first day, mice are trained for 5mins, starting at 6cm/s; the speed is slowly increased to 10cm/s. The treadmill has a 5° inclination and delivers a 0.2mA shock. The following day, measurement is performed: initial speed is set up at 10cm/s, increased of 2cm/s every minute. The test ends at exhaustion, when the mouse spends more than three seconds on the electric shock grid. Distance covered, speed and number of shocks were recorded.

### Rotarod

*Mpz*R98C/+ mice were evaluated at PND120 and PND180 on an accelerating rotating rod. Mice underwent three training trials on an IITC Life Science Roto-Rod (Series 8) with a ramp speed from 2 to 20rpm in 300s. A 1h rest was given after each trial and it was considered valid if the animals ran forward on the rod for at least 10s. The next day the latency to fall was recorded three times following the above-mentioned protocol for each time point and mouse. The average was used as the outcome value.

### Grip strength

#### C3-PMP22 study

The muscular strength was evaluated using a GSM Grip-Strength Meter (Ugo basile). This test measures the muscular strength using an isometric dynamometer connected to a grid. Once the animal is holding the grid with its forepaws it is slowly moved backwards pulling the tail, until it releases the grip. The dynamometer records the maximal force exerted. Each mouse is tested six times.

#### *Mpz*R98C/+ study

The strength of all four limbs of WT and *Mpz*R98C/+ mice was evaluated using an automated Grip Strength Meter (Columbus Instruments). Within 1 week after training (10 practice trials using a mesh bar), the peak force exerted by each individual mouse was measured 10 times consecutively with 10s resting periods and averaged.

### Electrophysiological analysis

#### C3-PMP22 study

The electrophysiological evaluation was performed with a specific EMG system (NeuroMep Micro, Neurosoft, Russia), as previously described [5]. Mice were anesthetized with trichloroethanol, 0.02ml/g of body weight, and placed under a heating lamp to maintain constant body temperature. Sciatic nerve conduction velocity was obtained by stimulating the nerve with steel monopolar needle electrodes. A pair of stimulating electrodes was inserted subcutaneously near the nerve at the ankle. A second pair of electrodes was placed at the sciatic notch to obtain two distinct sites of stimulation, proximal and distal along the nerve. Compound motor action potential (CMAP), was recorded with a pair of needle electrodes; the active electrode was inserted in muscles in the middle of the paw, whereas the reference was placed in the skin between the first and second digit. Sciatic nerve F-wave latency measurement was obtained by stimulating the nerve at the ankle and recording the responses in the paw muscle, with the same electrodes employed for the NCV study.

#### *Mpz*R98C/+ study

Mice were analyzed as above with the following differences. Anesthesia was with Ketamine/Xylazine (87.5mg/kg ketamine, 12.5mg/kg Xylazine). For sensory electrophysiological testing, one pair of loop electrodes were put 0.2cm and 0.7cm from the tail base as recording and reference electrodes. A second air was placed on the tail 3.7cm (cathode) and 4.2cm (anode) from the base. A ground electrode was put at the middle of two pairs loop electrodes. The distal pair was used for stimulation.

### Morphological analysis

Mice were sacrificed at the indicated time points and sciatic and femoral nerves were dissected. Semi-thin section and electron microscope analyses were performed as previously described [14]. The number of amyelinated axons in C3-PMP22 mice was counted blind to genotype and treatment from quadricep femoral nerve semi-thin sections (0.5-1μm thick) stained with toluidine blue, on images taken with a 100x objective, after whole nerve reconstruction. Similarly, for *Mpz*R98C/+ mice, semithin sections were examined under a 63x objective. Each consecutive field was captured using a digital camera and whole nerve reconstruction was made by using photoshop software. In both studies, g-ratio analysis (axonal diameter/fiber diameter) and the size distribution of myelinated fibers (based on axons diameter) were also measured for all fibers. 4-8 mice per genotype and condition were analysed. Ultrathin sections (90nm thick) from femoral nerves were cut using an ultracut ultramicrotome, stained with uranyl acetate and lead citrate and examined by transmission electron microscopy (TEM) (Zeiss Leo912 Omega or JEOL 1230, Peabody, MA).

### Protein extraction and Western Blotting

Mice were sacrificed at the indicated time points and sciatic nerves were dissected and immediately frozen in liquid nitrogen. Protein extraction was performed as previously described [52, 71]. The following antibodies were used: rabbit anti-Grp78/Bip (1:1000, Novus Biological, NB300-520 or 1:1000, Abcam, ab21685), rabbit anti-Phospho-eIF2α (Ser51) (D9G8) XP™ and eIF2α (D7D3) XP™ (1:2000, Cell Signalling, #3398 and #5324); chicken anti-P0 (PZO, Aves); rabbit anti-PMP22 (AB211052 or AB 861220; ABCAM); mouse anti-β-tubulin (1:1000, T4026, Sigma); mouse anti-Gapdh (1:1000, Millipore, MAB374); rabbit anti c-Jun (1:1000, ABCAM, #32137). Peroxidase-conjugated secondary antibodies (anti-rabbit HRP, DAKO, P0448; anti-chicken IgG-peroxidase, Sigma) were visualized using Amersham ECL or ECL Prime 225 reagent (GE Healthcare) for high-sensitivity chemiluminescent protein detection with UVItec gel analysis systems or using enhanced chemiluminescence (ECL) reagents (Bio-Rad) with autoradiography film (Kodak Scientific Imaging Film, Blue XB). Total proteins were visualized via staining with Coomassie Brilliant blue R250 staining solution (Bio-Rad). Densitometric quantification was performed with ImageJ.

### RNA isolation and Real-time PCR analysis

#### C3-PMP22 study

Total RNA was extracted using TRIzol (Roche Diagnostic GmbH, Germany) and reverse transcription was performed as described previously [10] Quantitative RT-PCR was performed according to manufacturer’s instructions (TaqMan, PE Applied Biosystems Instruments) on an ABI PRISM 7700 sequence detection system (Applied Biosystems Instruments). Normalization was performed using 18S rRNA as reference gene. Target and reference gene PCR amplification were performed in separate tubes with Assay on Demand™ (Applied Biosystems Instruments): 18S assay Hs99999901_s1; Ddit3/Chop, Mm00492097_m1; Xbp-1s assay, Mm03464496_m1; Hspa5/BiP assay, Mm00517691_m1; ATF4 assay, Mm00515324_m1; Gadd34/PPP1r15a assay, Mm00435119_m1.

#### *Mpz*R98C/+ study

Total RNA was extracted using NucleoSpin RNA Plus Kit (740984.50 Macherey-Nagel GmbH & Co. KG, Germany). Complementary DNA was prepared with SuperScript® III First Strand Synthesis SuperMix (11752-050, Invitrogen), and samples were analyzed as triplicates on a StepOnePlus Real-Time PCR System (Applied Biosystems) detection system using Fast SYBR® Green (4385612, Applied Biosystems). All samples were normalized to Ppia as an endogenous control and expressed relative (threshold cycle (Ct) and 2(-ΔΔCt)) to vehicle sciatic nerve data. The list of primers is below:

CHOP forward 5’-CTGCCTTTCACCTTGGAGAC-3’

CHOP Reverse 5’-CGTTTCCTGGGGATGAGATA-3’

BiP forward 5’-CATGGTTCTCACTAAAATGAAAGG-3’

BiP Reverse 5’-GCTGGTACAGTAACAACTG-3’

Xbp1-s forward 5’-GAGTCCGCAGCAGGTG-3’

Xbp1-s reverse 5’-GTGTCAGAGTCCATGGGA-3’

Ppia forward 5’-AGCACTGGAGAGAAAGGATT-3’

Ppia reverse 5’-ATTATGGCGTGTAAAGTCACCA-3’

### IFB-088 pharmacokinetic study

Nine males and nine females C57BL/6J (three animals per sampling time) were treated with a single intraperitoneal administration of IFB-088 at 4 mg/kg. 10min, 30min, 1h, 2h, 4h, 6h, 8h after administration, animals were briefly anaesthetized with Isofluorane and blood was collected. IFB-088 was extracted from plasma samples and measured by LC-MS/MS.

### Statistical analysis

Group sizes were pre-determined with the G*Power v3.1.9.4 software (Heinrich-Heine-Universität Düsseldorf), to detect differences of at least 1,25 standard deviations between groups, with 80% power an alpha error of 0.05. Data were analysed with GraphPad Prism7.02 software. For behaviour and electrophysiology, outliers have been identified using Grubbs’s test and data tested for normality and variance homogeneity. Statistical difference between mean values between two groups were tested using Student’s T-test or Mann-Whitney. When multiple groups were compared, one-way ANOVA analysis followed by Friedman’s, Tukey’s or Dunnett’s multiple comparison test and Kruskal-Wallis analysis followed by Dunn’s multiple comparison test were used, as indicated in the Figure legends.

## Results

### *Mpz*R98C/+ DRG Schwann cell co-cultures show improved myelination following treatment with IFB-088

Explanted dorsal root ganglia (DRGs) from wild type (WT) and *Mpz*R98C/+ mice were plated and grown under myelinating conditions [12]. Similar to what was found in explants from *Mpz*Ser63del mice [12] the percentage of myelinated internodes was reduced in explant cultures of *Mpz*R98C/+ mice compared to explants from WT animals (Additional File 1: Supplementary Fig. 1 a-b), although higher than in *Mpz*S63del co-cultures. *Mpz*R98C/+ co-cultures were then treated with escalating doses (50, 75, 100 and 125nM) of IFB-088 and assessed for myelination. The number of myelinated internodes increased with all doses compared to untreated co-cultures with a maximum effect at 100nM IFB-088 (Fig. 1a, Fig. 1b), similarly to what observed in *Mpz*S63del explants (Additional File 1: Supplementary Fig. 1c-d and [12]).

**Figure 1:**
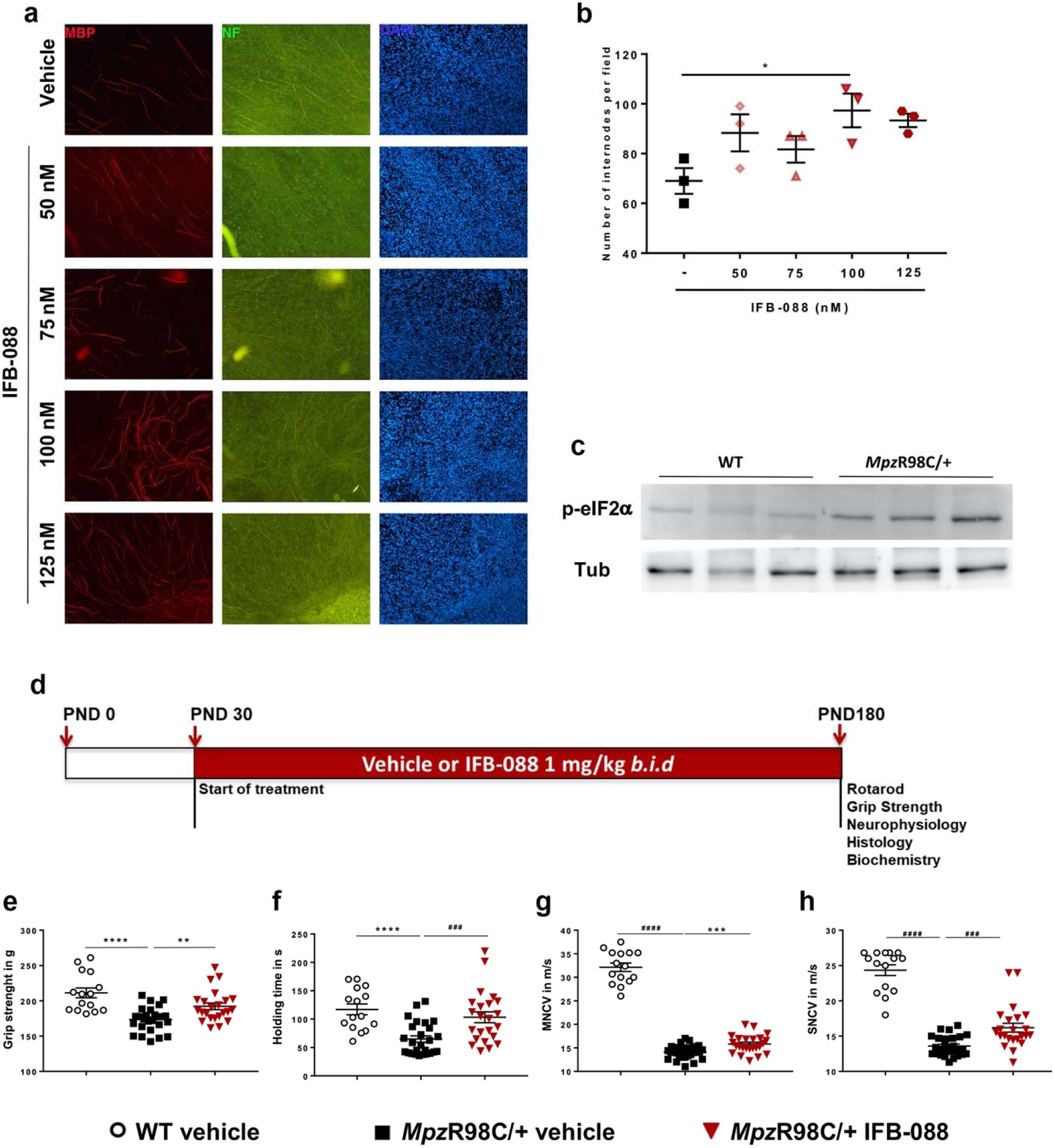
IFB-088 improves myelination in DRG explants, motor function and nerve conduction velocity from *Mpz*R98C/+ mice. Dorsal root ganglia (DRG) were dissected from embryos (E13.5) of *Mpz*R98C/+ mice. The myelination process was induced with ascorbic acid. After 2-week of treatment with vehicle or the indicated concentration of IFB-088, the DRGs were fixed and nuclei visualized by DAPI staining; axons and myelin were visualized by immunostaining with neurofilament antibody (NF) and with myelin basic protein (MBP) antibodies respectively. (**a**) Representative pictures. (**b**) Number of myelinated internodes per field in *Mpz*R98C/+ DRG explant cultures treated with vehicle or the indicated concentration of IFB-088 for 2 weeks. Mean ±SEM. *n*=3 independent experiments. \**P*<0.05 One-way ANOVA followed by Friedman’s test. (**c**) Representative western blot (WB) for P-eIF2_α_ and tubulin on sciatic nerve protein extracts from 1-month-old WT and *Mpz*R98C/+ mice. (**d**) Diagram of the treatment strategy. 30-day-old WT and *Mpz*R98C/+ mice were orally administered with vehicle or IFB-088 1mg/kg *b*.*i*.*d*. for 5-month. (**e**) Four limb grip strength max values average of 10 trials. Data were expressed in grams (g) as mean ±SEM. *n*=15-26 mice per condition. (**f**) Rotarod analysis. Data are expressed in seconds (s) as mean ±SEM. *n*=15-26 mice per condition. (**g**) Motor nerve conduction velocity (MNCV). Data are expressed in meter/second (m/s) as mean ±SEM. *n*=15-27 mice per condition. (**h**) Sensory nerve conduction velocity (SNCV). Data are expressed in meter/second (m/s) as mean ±SEM. *n*=15-26 mice per condition. \*\**P*<0.01; \*\*\**P*<0.001, \*\*\*\**P*<0.0001 by Student’s T-test; ###*P*<0.001, ####*P*<0.0001 by Mann-Whitney.

### IFB-088 treatment improves grip strength, rotarod performance and nerve conduction velocities in *Mpz*R98C/+ mice

Western blot (WB) on sciatic nerve lysates from 1-month old *Mpz*R98C/+ mice, which already manifested signs of the disease [56], showed a clear increase of P-eIF2α (Fig. 1c), indicating activation of the PERK pathway of the UPR. We treated 1-month old WT and *Mpz*R98C/+ mice with either vehicle or 1mg/kg of IFB-088 twice a day (the dosage shown to be effective in *Mpz*S63del mice) [12] by oral gavage. Throughout the treatment mice were regularly weighted. As previously reported for *Mpz*S63del mice [12], IFB-088 treatment does not impact body weight (Additional File 1: Supplementary Fig. 2).

After 3- and 5-month of treatment mice were tested by Rotarod, Grip Strength, and by neurophysiology. The mice were then sacrificed for morphological and biochemical analysis (Fig. 1d and Additional File 1: Supplementary Fig. 3a). As previously observed [12], IFB-088 treatment did not impact WT mice motor function or neurophysiological parameters (not shown). *Mpz*R98C/+ mice had reduced grip strength compared to WT animals at PND120 (Additional File 1: Supplementary Fig. 3b) and PND180 (Fig. 1e). After 5-month treatment *Mpz*R98C/+ mice showed significant improvement on grip strength compared to vehicle treated animals (Fig. 1e). Similar improvements were obtained for male and female mice (not shown). We evaluated the mice on the same day for their ability to maintain balance on an accelerating Rotarod. We confirmed a significant reduction in the latency to fall in PND120 and PND180 *Mpz*R98C/+ mice as compared to WT mice (Additional File 1: Supplementary Fig. 3c and Fig. 1f) [52, 56]. *Mpz*R98C/+ mice treated with IFB-088 for 5 months, but not 3 months, were able to maintain their balance significantly longer than untreated animals, approaching latencies obtained by WT animals (Fig. 1f and Additional File 1: Supplementary Fig. 3c).

Neurophysiological testing confirmed marked slowing in motor nerve conduction velocity (MNCV) and sensory nerve conduction velocity (SNCV) in *Mpz*R98C/+ mice at PND120 (Additional File 1: Supplementary Fig. 3d-e) and PND180 (Fig. 1g-h) [52, 56]. No significant differences in compound muscle action potential (CMAP) and sensory nerve action potential (SNAP) amplitudes were observed between WT and *Mpz*R98C/+ mice (not shown). 5-month treatment with IFB-088 significantly improved both MNCV and SNCV of *Mpz*R98C/+ mice (Fig. 1g-h). The positive impact of IFB-088 treatment on *Mpz*R98C/+ SNCV was also observed after 3-month of treatment (Additional File 1: Supplementary Fig. 3e).

### IFB-088 treatment improves nerve morphology, partially reduces ER-stress and corrects the expression of Schwann cell differentiation marker in *Mpz*R98C/+ mice

On morphometric analysis, nerves from *Mpz*R98C/+ mice demonstrated marked abnormal myelin as attested by reduced myelin thickness and increased g-ratio (Fig. 2). After 5-month IFB-088 treatment, transverse sections of quadriceps femoral nerves from *Mpz*R98C/+ mice showed increased myelin thickness and reduced g-ratio (ratio between axon diameter and axon + myelin diameter) compared to vehicle treated nerves (Fig. 2a-e). Detailed ultrastructural analysis performed via EM, confirmed the thin myelin sheath surrounding *Mpz*R98C/+ axons, which appears thicker after IFB-088 treatment (Fig. 2f).

**Figure 2:**
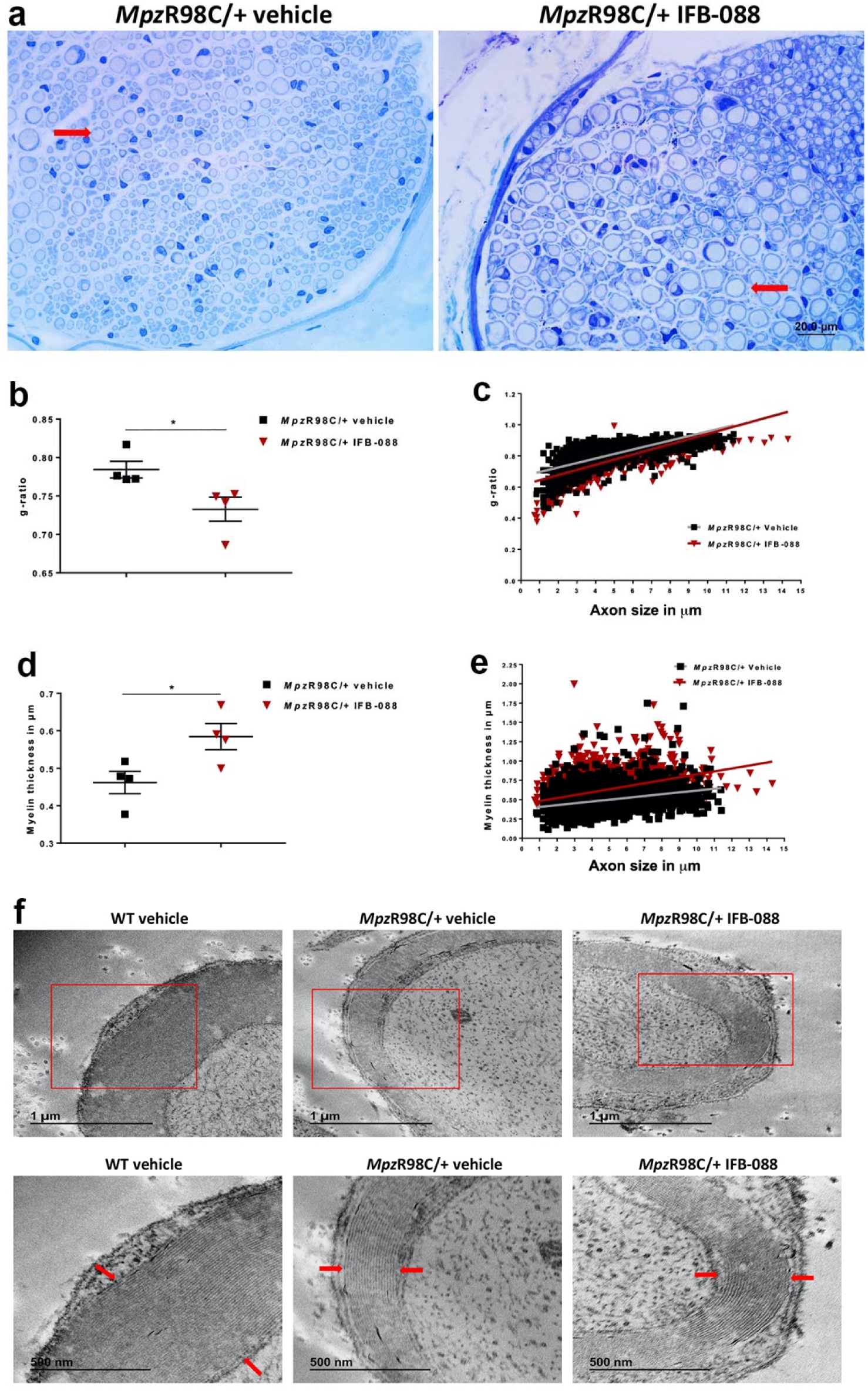
IFB-088 treatment improves *Mpz*R98C/+ mice quadriceps femoral nerve morphology. **(a)** Toluidine blue stained semithin sections of quadriceps femoral nerve from *Mpz*R98C/+ mice treated with vehicle *b*.*i*.*d*. or IFB-088 1mg/kg *b*.*i*.*d*. for 5 months. Arrows illustrate myelinated axons. Scale bar, 20µm. (**b**) Quadriceps femoral nerve g-ratios from *Mpz*R98C/+ mice treated with vehicle *b*.*i*.*d*. or IFB-088 1mg/kg *b*.*i*.*d*. for 5 months. Data are expressed as mean ±SEM. *n*=4 mice per condition. \**P*<0.05 by Student’s T-test. (**c**) Scatter plot of quadriceps femoral nerve g-ratios toward axon diameter from *Mpz*R98C/+ mice treated with vehicle *b*.*i*.*d*. or IFB-088 1mg/kg *b*.*i*.*d. n*=4 mice per condition. (**d**) Myelin thickness of quadriceps femoral nerve from *Mpz*R98C/+ mice treated with vehicle *b*.*i*.*d*. or IFB-088 1mg/kg *b*.*i*.*d*. for 5 months. Data are expressed in µm as mean ±SEM. *n*=4 mice per condition. \**P*<0.05 by Student’s T-test. (**e**) Scatter plot of quadriceps femoral nerve myelin thickness toward axon diameter from *Mpz*R98C/+ mice treated with vehicle *b*.*i*.*d*. or IFB-088 1mg/kg *b*.*i*.*d. n*=4 mice per condition. (**f**) Ultrathin sections of quadriceps femoral nerve from WT mice treated with vehicle *b*.*i*.*d*. and from *Mpz*R98C/+ mice treated with vehicle *b*.*i*.*d*. or IFB-088 1mg/kg *b*.*i*.*d*. for 5 months. Lower panels are higher magnification images of upper panels. Scale bar 1µm for top panels, and 500nm for bottom panels.

We previously showed that *Mpz*R98C/+ mice have increased expression of ER-chaperones immunoglobulin heavy chain-binding protein (BiP) and glucose regulated protein 94 (Grp94), as part of an ER-stress response. We also demonstrated an altered phenotype of myelinating Schwann cells in which the expression of c-Jun, a transcription factor that inhibits myelination [50], was increased [56]. Therefore, we investigated the expression of mRNA (Fig. 3a-c), and protein levels (Fig. 3d-f) in a variety of genes known to play a role in these processes. As previously described, *Mpz*R98C/+ mice presented an increase of the expression of ER-stress and UPR markers *BiP*, C/EBP homologous protein (*Chop*) and spliced X-box-binding protein-1 (*Xbp1s*) and a higher level of c-Jun compared to WT mice (Fig. 3). IFB-088 treatment reduced the levels of *Chop* and *BiP* mRNA and BiP protein, whereas the levels of *Xbp1s* were unchanged (Fig. 3a-e). Finally, c-Jun protein levels were significantly reduced by IFB-088 (Fig. 3d, Fig. 3f). Altogether our data suggest that treatment with IFB-088 reduced stress levels in *Mpz*R98C/+ nerves, allowing Schwann cell to differentiate and myelinate more efficiently.

**Figure 3:**
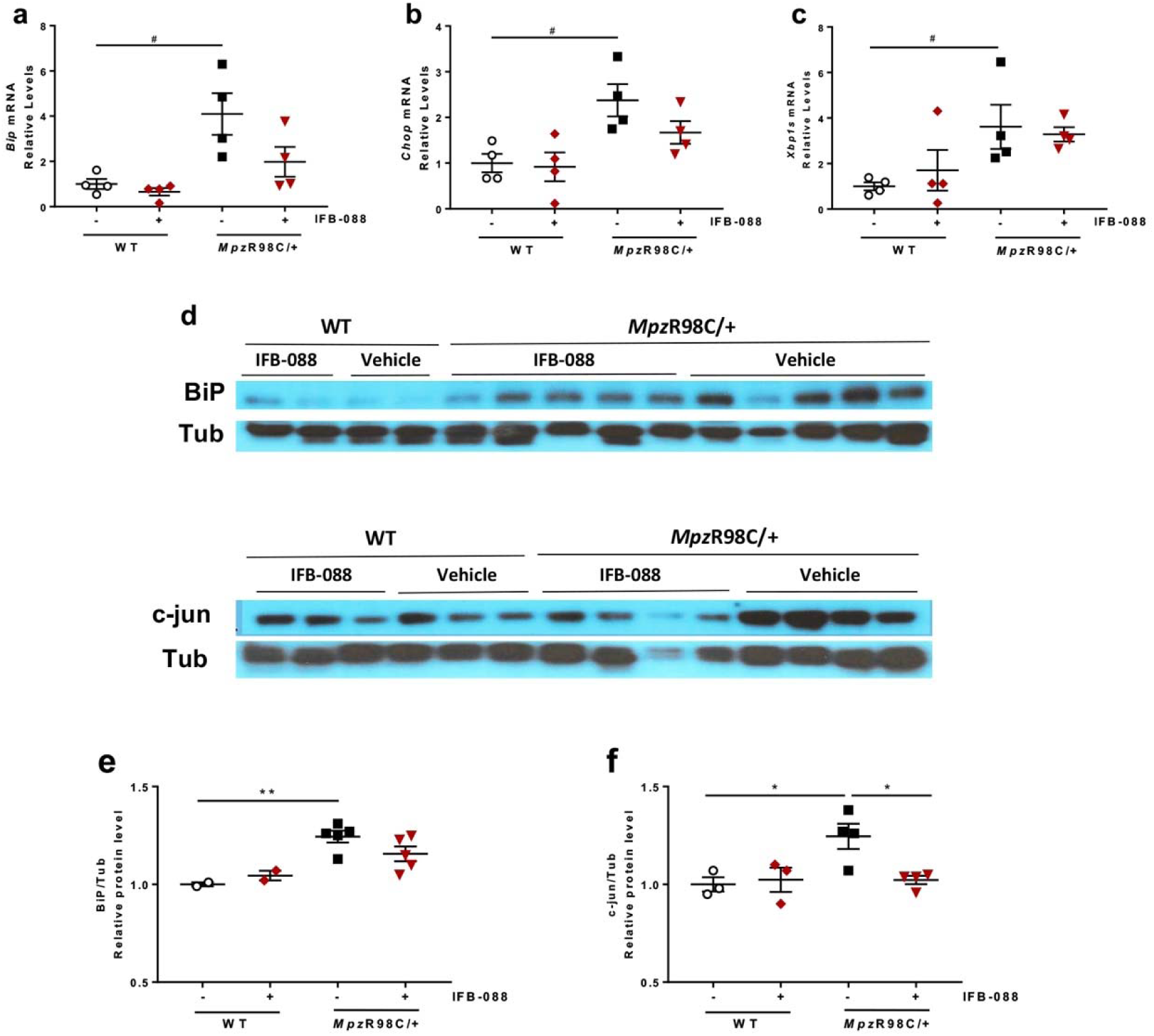
IFB-088 treatment reduces ER stress and Schwann cells differentiation markers expression in *Mpz*R98C/+ sciatic nerve. Evaluation of mRNA (**a-c**) and protein levels (**d-f**) on sciatic nerve samples from WT and *Mpz*R98C/+ mice treated with vehicle *b*.*i*.*d*. or IFB-088 1mg/kg *b*.*i*.*d*. for 5 months. mRNA relative levels of *Bip* (**a**), *Chop* (**b**) and *Xbp1s* (**c**) determined by qRT-PCR. *n*=4 per condition. #*P*<0.05 by Mann-Whitney. (**d**) Western blot images for BiP and c-Jun. Quantification relative to tubulin for BiP (**e**), and c-Jun (**f**). *n*=2-5 per condition. \**P*<0.05, \*\**P*<0.01 by Student’s T-test.

### Nerves from the CMT1A mouse model C3-PMPP22 show alteration of myelin proteins stoichiometry and activation of ER-stress pathways

To address whether IFB-088 could be a viable treatment also for CMT1A we took advantage of the C3-PMP22 mouse (supposed to carry 3-4 extra copies of PMP22) [65]. WB performed of protein extract from C3-PMP22 nerves at 4-month demonstrated reduced expression of P0 (MPZ), the most abundant myelin protein, but similar PMP22 levels to WT nerves. As a result, there was a relative overexpression of PMP22 protein in C3-PMP22 nerves (Fig. 4a).

**Figure 4:**
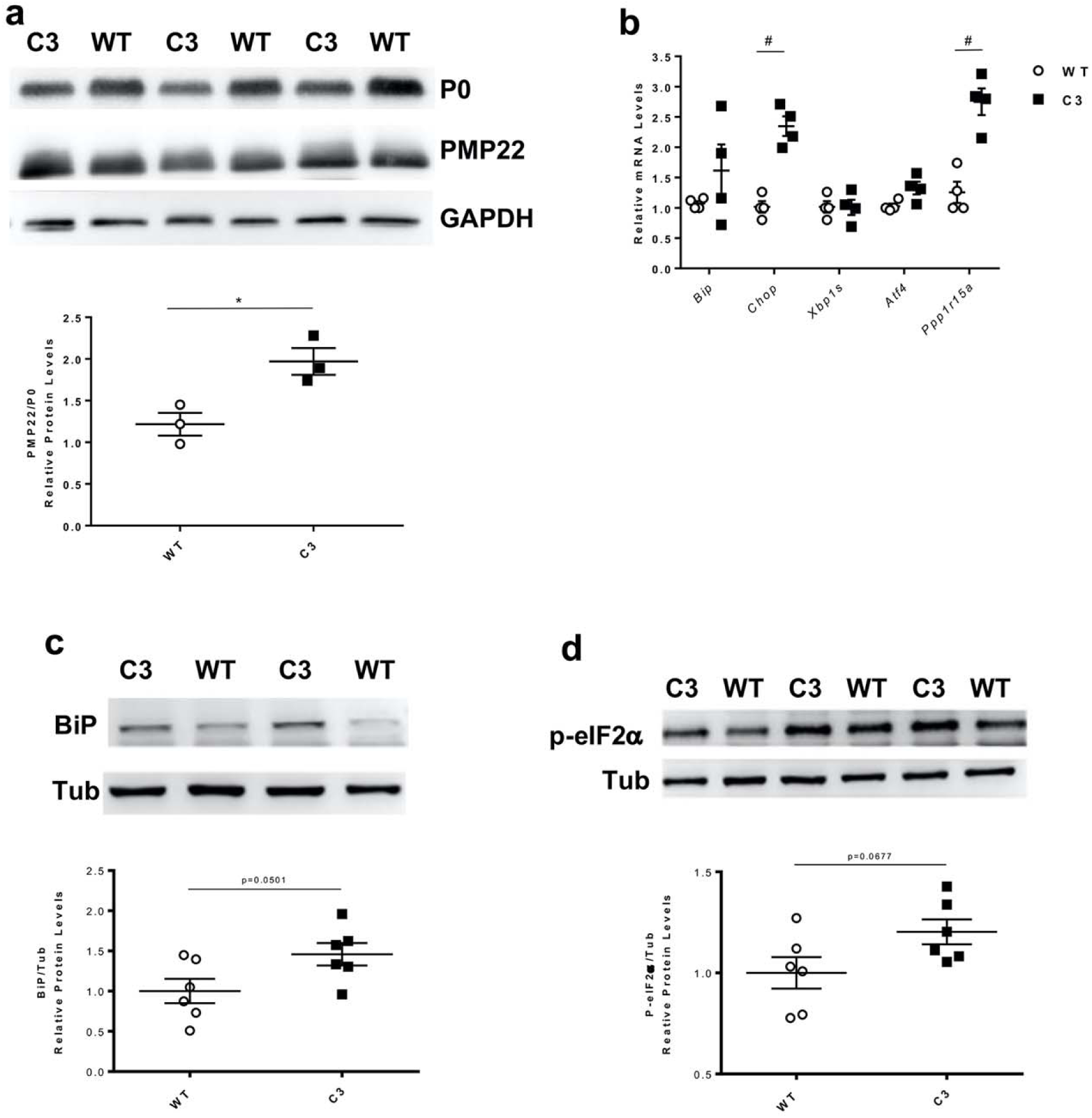
C3-PMP22 mice show a relative overexpression of PMP22 associated with expression of ER stress/UPR markers. (**a**) Evaluation of PMP22, P0 and GAPDH protein levels by WB in sciatic nerve protein lysates from 4-month-old WT and C3-PMP22 (C3) mice. Top: Representative picture. Bottom: Quantification of the PMP22/P0 protein ratio. *n*=3 per condition. \**P*<0.05 by Student’s T-test. (**b**) Evaluation of *Bip, Chop, Xbp1s, Atf4* and *Gadd34/Ppp1r15a* mRNA levels by qRT-PCR on sciatic nerve samples from 4-month-old WT and C3-PMP22 (C3) mice. *n*=4 per condition. #*P*<0.05 Mann Whitney. (**c**) Evaluation of BiP and tubulin protein levels by WB in sciatic nerve protein lysates from 4-month-old WT and C3-PMP22 (C3) mice. Top: Representative picture. Bottom: Quantification relative to tubulin. *n*=6 per condition. Student’s T-test. (**d**) Evaluation of eIF2_α_ phosphorylation level and tubulin protein level by WB in sciatic nerve protein lysates from 1-month-old WT and C3-PMP22 (C3) mice. Top: Representative picture. Bottom: quantification of eiF2_α_ phosphorylation level relative to tubulin. *n*=6 per condition. Student’s T-test.

To test whether the excess of PMP22 would lead to proteostatic stress we measured the mRNA levels for a subset of well know UPR markers. This revealed a small increase in the expression of the general ER-stress marker *BiP* (downstream of the ATF6 arm of the UPR), and a significant upregulation of *Chop* and of the Protein Phosphatase 1 Regulatory Subunit 15A (*Gadd34/Ppp1r15a*), two downstream targets of the PERK/P-eIF2α pathway, whereas *Xbp1s*, a target of the inositol requiring enzyme 1 (IRE1) arm, and the activating transcription factor 4 (*Atf4*) were not increased (Fig. 4b). WB for BiP and P-eIF2α confirmed the activation of stress pathways in these nerves (Fig. 4c-d). These results showed that C3-PMP22 mice present an alteration of myelin proteins stoichiometry associated with the activation of ER-stress pathways.

### IFB-088 treatment improves motor capacity and MNCV in CMT1A mice

We treated C3-PMP22 mice with either 0.5mg/kg or with 1mg/kg of IFB-088 twice a day by oral gavage. The treatment started at PND15, when the disease is already manifested as shown by severe myelination defects, altered myelin proteins stoichiometry and activation of ER-stress pathways (Additional File 1: Supplementary Fig. 4).

After 10-week of treatment C3-PMP22 mice were tested for motor capacity and, after 12-week of treatment, for neurophysiology. They were then sacrificed for morphological and biochemical analysis (Fig. 5a). Treadmill analysis showed that C3-PMP22 mice were severely impaired compared to WT mice (Fig. 5b). IFB-088 treatment at 1mg/kg *b*.*i*.*d*. showed a small but significant improvement as compared to vehicle treated mice (Fig. 5b). However, the operator (blind to treatment) noticed that all C3-PMP22 male mice ran very poorly. Therefore, the mice were stratified by gender. C3-PMP22 female mice treated with IFB-088 1mg/kg *b*.*i*.*d*. had a significant amelioration in motor capacity, that increased 2-fold compared to vehicle treated mice (Fig. 5c). Male mice also had a comparable 2-fold improvement with the same dosage, but the baseline was very poor (Fig. 5d).

**Figure 5:**
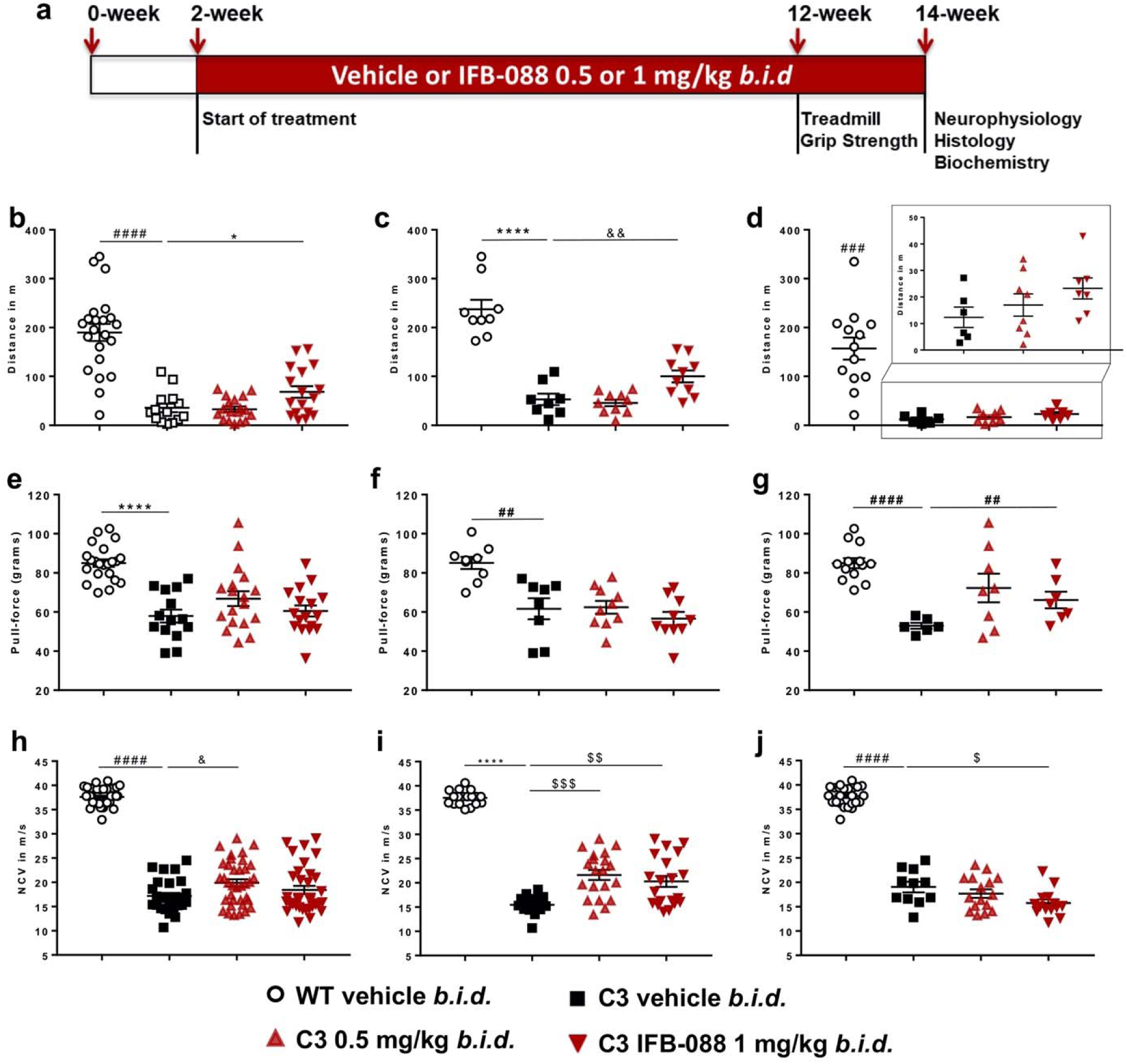
IFB-088 treatment improves motor function and nerve conduction velocity in C3-PMP22 mice. (**a**) Diagram of the treatment strategy. 15-day-old WT and C3-PMP22 (C3) mice were orally administered with vehicle *b*.*i*.*d*. or IFB-088 at 0.5 or 1mg/kg *b*.*i*.*d*. for 12 weeks. (**b**) Treadmill analysis performed after 10-week of treatment. Data from males and females expressed in meter (m) as mean ± SEM. *n*=14-22 mice per condition. (**c**) Data from females. *n*=8-10 mice per condition. (**d**) Data from males. *n*=6-13 mice per condition. (**e**) Forepaws grip strength average of 6 trials performed after 10-week of treatment. Data were expressed in pull force (grams) as mean ± SEM. *n*=14-22 mice per condition. (**f**) Data from females. *n*=8-10 mice per condition. (**g**) Data from males. *n*=6-13 mice per condition. (**h**) Motor nerve conduction velocity (MNCV) performed after 12-week of treatment. Data from males and females expressed in meter/second (m/s) as mean ±SEM. n=14-21 mice per condition. (**i**) Data from females. *n*=8-10 mice per condition. (**j**) Data from males. *n*=6-12 mice per condition. \**P*<0.05; \*\**P*<0.01; \*\*\*\**P*<0.0001 by Student’s T-test. #*P*<0.05; ###*P*<0.001; ####*P*<0.0001 by Mann-Whitney. &*P*<0.05; &&*P*<0.01 by One-Way ANOVA followed by Dunnett’s test. $*P*<0.05; $$*P*<0.01; $$$*P*<0.001 by Kruskal-Wallis followed by Dunn’s test.

Grip strength confirmed that C3-PMP22 mice were impaired compared to WT mice. IFB-088 treatment did not improve strength when the mice were analysed altogether (Fig. 5e). However, when divided by gender, C3-PMP22 male mice showed a trend towards improvement at 0.5mg/kg *b*.*i*.*d*. which became significant for the 1mg/kg *b*.*i*.*d*. of IFB-088 (Fig. 5g). This positive impact was not seen in C3-PMP22 female mice (Fig. 5f).

Finally, we assessed a series of neurophysiological parameters (MNCV, F-wave latency, CMAP) which revealed a severe impairment in C3-PMP22 mice as compared to WT controls (Fig. 5h and not shown), as previously reported [65]. Treatment with both 0.5 and 1mg/kg *b*.*i*.*d*. IFB-088 showed a modest improvement in MNCV after 12-week treatment, that reached significance for the 0.5mg/kg *b*.*i*.*d*. dose (Fig. 5h). Stratification per gender revealed that the improvement was restricted to female mice, for which the amelioration was statistically significant at both dosages (Fig. 5i). Only IFB-088 treated C3-PMP22 female mice displayed MNCV over 20m/s. On the contrary in C3-PMP22 male mice there was no improvement, and actually the 1mg/kg *b*.*i*.*d*. dose showed a small worsening that, although marginal, reached statistical significance (Fig. 5j). None of the other neurophysiological parameters measured were improved by the treatment (not shown).

### IFB-088 treatment improves nerve morphology, reduces ER-stress level and partially readjust myelin proteins stoichiometry in CMT1A mice

Previous analysis suggested that C3-PMP22 mice motor nerves were more affected than sensory nerves [24, 65]. Therefore, we dissected quadriceps femoral nerve, predominantly motor, and sciatic nerve which is a mixed nerve with many sensory axons. Transverse sections of sciatic nerves showed an increase in the myelination of large calibre fibres in mice treated with IFB-088. In both male and female C3-PMP22 nerves large calibre axons were almost invariably amyelinated (where by amyelinated we refer to any axon with diameter>1µM, which is in contact with a Schwann cell but not myelinated by it). After IFB-088 treatment many of the large axons were myelinated (Additional File 1: Supplementary Fig. 5). Similarly, in untreated femoral nerves most large calibre axons were amyelinated whereas small calibre axons tended to be hypermyelinated (Fig. 6a-b). As in sciatic nerves, treatment with IFB-088 partially corrected these abnormalities with myelin wrapping appearing around large axons at both 0.5 and 1mg/kg *b*.*i*.*d*. of IFB-088 (Fig. 6a-b). No gross differences were detected between female and male nerves (Additional File 1: Supplementary Fig. 5 and Fig. 6a-b).

**Figure 6:**
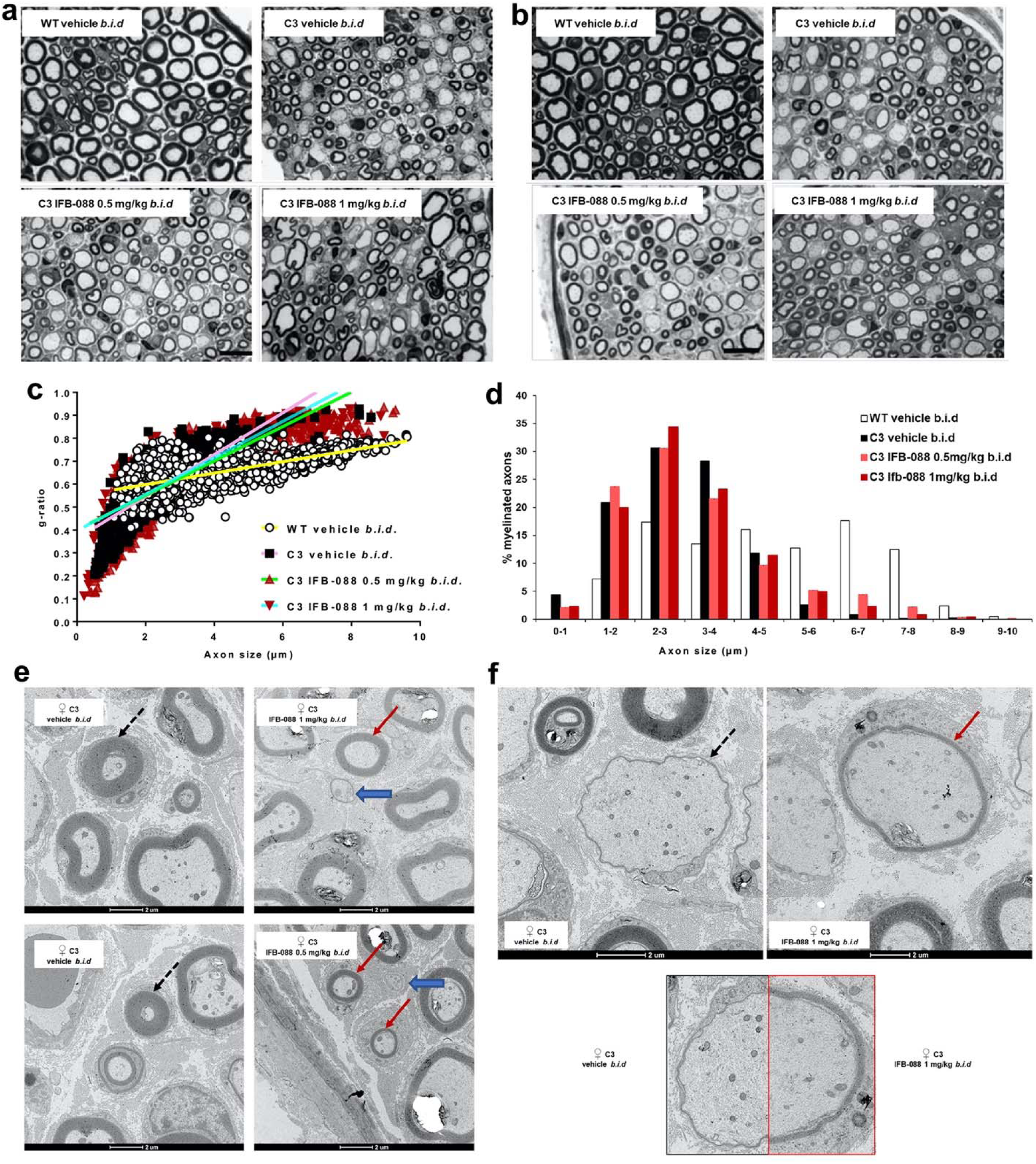
IFB-088 treatment improves C3-PMP22 mice quadriceps femoral nerve morphology. Toluidine blue stained semithin sections of quadriceps femoral nerve from (**a**) female and (**b**) male WT mice treated with vehicle *b*.*i*.*d*. and C3-PMP22 (C3) mice treated with vehicle *b*.*i*.*d*. or IFB-088 at 0.5 or 1mg/kg *b*.*i*.*d*. for 12 weeks. Scale bar, 10µm. (**c**) Scatter plot of quadriceps femoral nerve g-ratios from WT mice treated with vehicle *b*.*i*.*d*. and C3-PMP22 (C3) mice treated with vehicle *b*.*i*.*d*. or IFB-088 at 0.5 or 1mg/kg *b*.*i*.*d*. Note the “cloud” of axons with diameter lower than 1µm present only in C3-PMP22 nerves (untreated and treated) and the appearance of a considerable number of myelinated axons larger than 5-6µm in IFB-088 treated C3-PMP22 nerves. *n*=5-8 nerves per condition. (**d**) Percentage of myelinated axons per axons size. *n*=5-8 nerves per condition. (**e** and **f**) TEM analysis of quadriceps femoral nerve from C3-PMP22 (C3) mice treated with vehicle *b*.*i*.*d*. or IFB-088 at 1mg/kg *b*.*i*.*d*. (**e**) In C3-PMP22 vehicle treated nerves, small calibre axons are abnormally hypermyelinated (black dotted arrows). Treatment with IFB-088 results in normal looking myelin in axons larger than 1µm (red arrows) or in almost no myelin is smaller axons (blue larges arrows). (**f**) In C3-PMP22 vehicle treated nerves large calibre axons are basically amyelinated or with only a very thin layer of myelin (left panel). After treatment with IFB-088 (right panel), a subgroup of large calibre axons showed a properly compacted (albeit still rather thin) myelin sheath. A side by side comparison of myelination in two axons with similar calibre is shown in the lower panel.

Cross sections of the entire femoral nerve from 2-4 mice per condition for both genders were reconstructed and, for each nerve, the entire number of myelinated or amyelinated axons was counted (Table 1).

**Table 1.** Percentage of amyelinated (amongst axons with diameter larger than 1 µm) and myelinated fibres. Data were expressed in percentage of fibers as mean ± SEM. *n*=5-8 mice per condition. \**P*<0.05, \*\**P*<0.01 by One-way ANOVA with Tukey post-hoc test; §*P*<0.05 by Student’s T-test vs C3-PMP22 vehicle.

|  |  | WT Vehicle<br>*b.i.d.* | C3 Vehicle<br>*b.i.d.* | C3 IFB-088<br>0.5 mg/kg *b.i.d.* | C3 IFB-088<br>1 mg/kg *b.i.d.* |
| --- | --- | --- | --- | --- | --- |
| % of amyelinated<br>fibers | All mice | 0.033 ± 0.08 | 27.2 ± 2.4 | **19.9 ± 1.3\*\*** | 21.9 ± 1.4 |
|  | Females | 0.1 ± 0.06 | 29.9 ± 1 | 19.8 ± 2.5§ | 23.5 ± 0.3 |
|  | Males | 0 | 25.4 ± 3.9 | 20 ± 1.2 | 20.3 ± 2.7 |
| % of myelinated<br>fibers | All mice | 100 ± 0.026 | 72.8 ± 2.4 | **80 ± 1.3\*** | 78.1 ± 1.4 |
|  | Females | 99.8 ± 0.061 | 70.1 ± 1 | 80.1 ± 2.5§ | 76.5 ± 0.3 |
|  | Males | 100 | 74.6 ± 3.9 | 80 ± 1.2 | 79.7 ± 2.7 |

This analysis identified 27.2 ± 2.4% amyelinated large axons in C3-PMP22 vehicle treated nerves. Treatment with IFB-088 0.5 or 1mg/kg *b*.*i*.*d*. reduced the percentages to 19.9±1.3% and 21.9±1.4%, representing a 27% and 20% improvement respectively. The number of amyelinated axons was slightly greater for females (29.9±1%) than for males (25.4±3.9%) which also showed more variability. In males the treatment with IFB-088 0.5 or 1mg/kg *b*.*i*.*d*. had a similar effect, reducing amyelinated axons by roughly 20%, whereas in females the IFB-088 0.5mg/kg *b*.*i*.*d*. treatment reduced amyelinated axons by 34%, while the IFB-088 1mg/kg *b*.*i*.*d*. treatment reduced it by 20% (Table1).

Analysis of myelin thickness by g-ratio showed a remarkable difference in the scatter plot between WT and C3-PMP22 mice. The slope for C3-PMP22 mice was much steeper, mostly due to the large group of hypermyelinated small axons (diameter < 1 µM) accompanied by the almost complete absence of myelination in axons larger than 5µM (Fig. 6c). Indeed, whereas there were no myelinated axons smaller than 1µM in WT nerves, 4.4% of the axons myelinated in quadriceps femoral nerves from C3-PMP22 mice had a diameter lower than 1µM, indicating an aberrant hypermyelinating phenotype. Conversely, 45.9% of myelinated axons in nerves from WT mice had a diameter larger than 5µM, a percentage that was reduced to only 3.9% of the myelinated axons in C3-PMP22 mice (Fig. 6d). IFB-088 treatment partially corrected these abnormalities. The percentage of myelinated axons smaller than 1µM was reduced to 2.1% and 2.3% by the IFB-088 0.5mg/kg *b*.*i*.*d*. and 1mg/kg *b*.*i*.*d*. treatment respectively, whereas the percentage of myelinated axons with a diameter larger than 5µM rose to 12.4% and 8.5% respectively (Fig. 6d). Stratification of male and female mice showed that the amelioration in the hypermyelinating phenotype in small calibre axons was exclusively present in females, as visible in the scatterplots (Additional File 1: Supplementary Fig. 6a-b), even though the overall g-ratio in vehicle treated C3-PMP22 males and females was virtually identical (0.61±0.013 in males vs 0.61±0.015 in females). In both C3-PMP22 males and females, IFB-088 treatment increases the number of myelinated axons with diameter larger than 5µM (Additional File 1: Supplementary Fig. 6c-d).

Detailed analysis by transmission electron microscopy (TEM) on transverse sections from quadriceps femoral nerves confirmed that in C3-PMP22 nerves, small diameter axons (even smaller than 1_μ_m) were surrounded by an abnormally thick myelin sheath. In female nerves, treatment with IFB-088 almost completely corrected this phenotype (Fig. 6e and Additional File 1: Supplementary Fig. 6a). At the same time, large calibre axons were either not myelinated or presented a very thin layer of myelin (probably corresponding to 2-3 wraps of Schwann cell membrane). Treatment with IFB-088 allowed myelination to proceed also in large calibre fibres (Fig. 6f). Importantly, this newly formed myelin appeared to have a proper structure and compaction/periodicity.

As previously shown, C3-PMP22 nerves presented increase expression of the ER-stress marker BiP and an unbalance between PMP22 and P0 protein levels. We therefore wanted to test whether IFB-088 treatment could correct these features. WB analysis showed that treatment with both dosages of IFB-088 normalized BiP expression to WT levels (Fig. 7a-b), and that there was a partial (although not complete) readjustment of the PMP22/P0 protein ratio (Fig. 7a, Fig. 7c), suggesting an overall improvement in nerve proteostasis.

**Figure 7:**
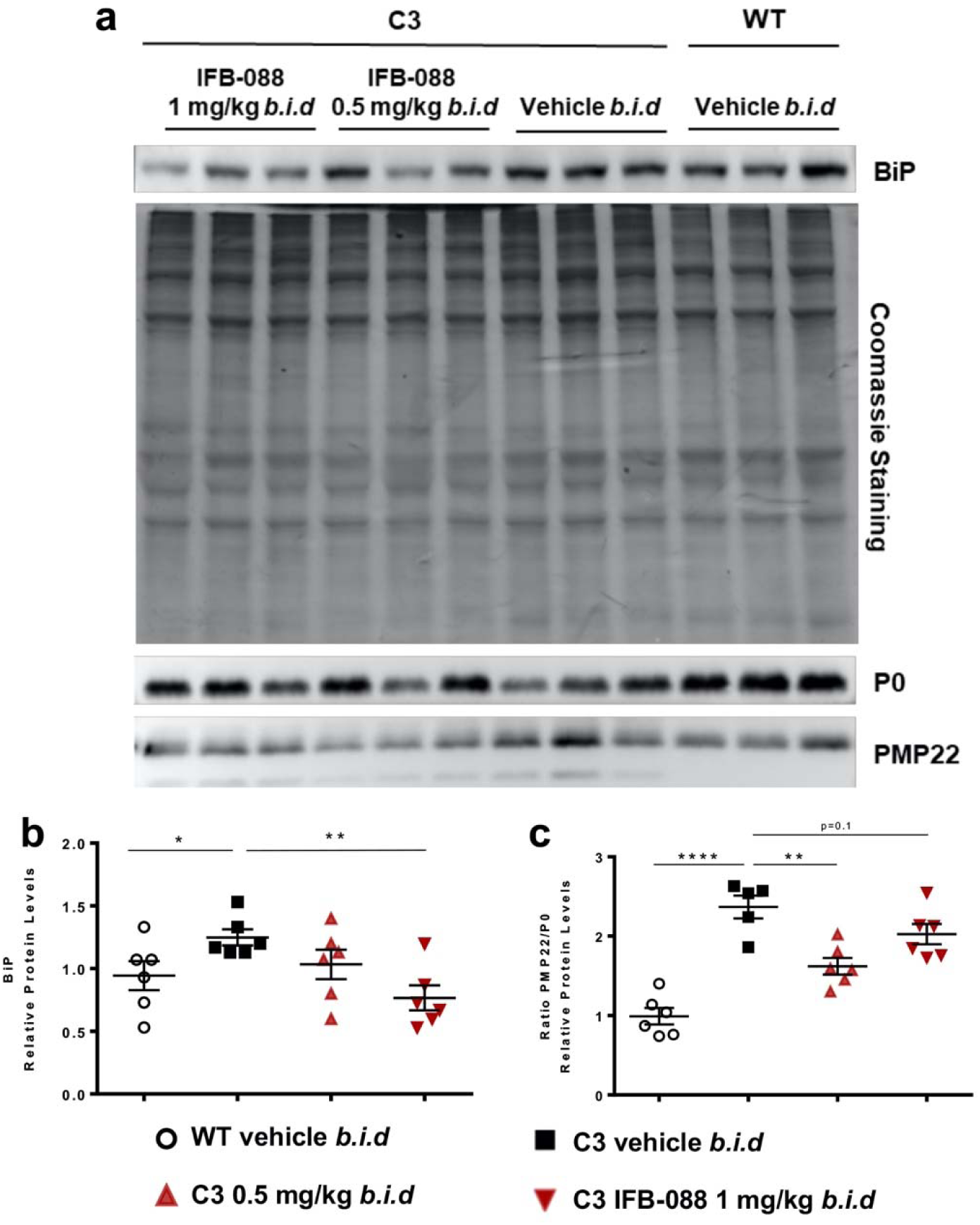
IFB-088 treatment improves PMP22 stoichiometry in C3-PMP22 mice peripheral nerves and reduces ER stress. Evaluation of PMP22, P0 and BiP protein levels by WB in sciatic nerve protein lysates from WT mice treated with vehicle *b*.*i*.*d*. and C3-PMP22 (C3) mice treated with vehicle *b*.*i*.*d*. or IFB-088 at 0.5 or 1mg/kg *b*.*i*.*d*. (**a**) Representative picture. Three representative samples per condition out of five/six are shown. (**b**) Quantification of BiP protein level. (**c**) Quantification of the PMP22/P0 protein ratio. \**P*<0.05; \*\**P*<0.01; \*\*\*\**P*<0.0001 by Student’s T-test.

## Discussion

The maintenance of correct protein homeostasis (proteostasis) is tightly controlled by protein quality control (PQC) mechanisms [1]. Myelinating Schwann cells, that need to synthesize large amounts of lipids and proteins, are particularly susceptible to failures in PQC. Alterations in proteostasis, deficits in PQC and the activation of ER-stress/UPR have been implicated in several myelin disorders [69]. Here we show that the small molecule IFB-088 is able to readjust protein homeostasis and to ameliorate disease features in two models of demyelinating CMT1, the *Mpz*R98C/+ (CMT1B) and the C3-PMP22 (CMT1A) mice.

### IFB-088 improves disease features in CMT1B mice

We demonstrated that IFB-088 effectively mitigates the peripheral neuropathy in *Mpz*R98C/+ mice, using *in vitro* as well as *in vivo* studies. *In vitro*, IFB-088 enhanced myelination in *Mpz*R98C/+ DRGs explant co-cultures. *In vivo*, 5-month IFB-088 treatment improved motor performance, nerve conduction velocity, and peripheral nerve morphology of *Mpz*R98C/+ mice. Our prior studies demonstrated that ER-chaperones such as BiP and the transcription factor CHOP are upregulated in *Mpz*R98C/+ mice consistent with their ER-stress response. We also previously demonstrated that the transcription factor c-Jun, which inhibits PNS myelination [50], was upregulated in *Mpz*R98C/+ mice [56]. We now show that treatment with IFB-088 reduced ER-stress and decreased the expression of the negative regulator of myelination c-Jun.

IFB-088 treatment effects, while significant in *Mpz*R98C/+ mice, were less pronounced than those observed in *Mpz*Ser63del animals [12]. Morphological as well as molecular deficits were often prevented in the *Mpz*Ser63del mice [12] but only showed improvement in the *Mpz*R98C/+. We posit several reasons for these differences. First, *Mpz*R98C/+ mice may be considered a “more authentic disease model” than the *Mpz*Ser63del animals, and this may contribute to the different responses to treatment between the two models. The *Mpz*R98C/+ mice have had the mutation “knocked in” to the endogenous mouse *Mpz* allele [56]. Alternatively, the *Mpz*Ser63del mice are transgenic animals and still retain two WT *Mpz* alleles in addition to the *Mpz*Ser63del expressing transgene [70]. Thus, *Mpz*Ser63del mice have two copies of WT *Mpz* as opposed to the one *Mpz*R98C/+ mice to generate the normal protein. Secondly, the R98C *MPZ* neuropathy in humans is more severe than that caused by Ser63del *MPZ*. Patients with R98C *MPZ* (also called R69C by an earlier numbering system) have a dysmyelinating neuropathy and present with delayed developmental milestones, often not walking independently until after 2 years of age. Affected patients may require walkers or wheelchairs for ambulation prior to reaching adulthood, and typically have MNCV<10 m/s [3, 55]. Patients with the Ser63del *MPZ* mutation typically reach developmental milestones on time, walk independently by a year of age and only slowly develop symptoms over the first two decades of life. They have not required more than ankle foot orthotics to ambulate even in adulthood [41]. In humans *MPZ*S63del MNCV are in the 20-30m/s range, similar to what is observed in patients with CMT1A [41]. Thus, clinically, the Ser63del *MPZ* mutation is considered to be a demyelinating rather than a dysmyelinating neuropathy. Taken together these data suggest the *Mpz*R98C mouse model causes a more severe neuropathy, that would be expected to be more difficult to reverse than the neuropathy caused by the Ser63del *Mpz* mutation.

How alleviating ER-stress improves the neuropathy caused by the R98C and Ser63del *MPZ* mutations remains an important issue. It is unlikely that augmenting UPR activity improves myelination by allowing R98C P0 to be transported to and inserted into PNS myelin. Introducing a cysteine into the P0 extracellular domain at codon 98 would be predicted to disrupt the disulfide bridge between existing cysteines at codons 50 and 127 [59]. This would in turn disrupt the secondary and tertiary structures of the extracellular domain that are necessary to create P0 tetramers in cis and trans to compact myelin [59], as already elegantly shown for the *Mpz*S63C mutation [2, 70]. We believe it more likely that treatment benefits from IFB-088 occur by facilitating the Schwann cells ability to degrade the R98C P0 as well as transport WT P0 to myelin. Myelinating Schwann cells generate very large amounts of proteins and lipids to form and maintain the myelin sheath [42, 44]. Processing and properly folding P0 glycoprotein in the ER is particularly demanding as P0 is, by far, the major PNS myelin protein, comprising approximately 50% of all PNS myelin proteins [22] and 2% of all Schwann cell transcripts during the peak of myelination [36]. Folding and post-translationally processing P0 by ERQC pathways is a major undertaking for the ER, even when the protein is in its WT form [13]. The task is increased when a mutation such as R98C makes folding the protein more difficult. Schwann cells typically target misfolded proteins for degradation, through ubiquitination and proteasomal processing. Both protein degradation and proteosome function have been shown to be impaired in *Mpz*Ser63del mice [67, 68] and this is likely to be the case with *Mpz*R98C/+ mice as well. We predict that IFB-088-mediated translational attenuation through persistent phosphorylation of eIF2α, will better enable Schwann cells to target R98C P0 to the proteosome, readjusting protein homeostasis as shown in *Mpz*Ser63del mice [11, 12]. In support of this hypothesis, it has been recently shown that promoting degradation by stimulating the proteasome ameliorates proteostasis and improves the phenotype of *Mpz*Ser63del mice [66]. We have previously shown that UPR activation causes *Mpz*R98C and *Mpz*S63del Schwann cells to enter a limited differentiation state in part by upregulating _C_-Jun expression; c-Jun, a transcription factor, negatively regulates myelination [50] [16]. As a result of this inhibition, myelin protein expression, including P0, decreases so that there is less mutant P0 for the ER to process [56].

IFB-088 treatment reduces the expression of the UPR marker BiP in *Mpz*R98C/+ mice, suggesting a reduction of ER-stress upon treatment. In addition, IFB-088 treatment decreases c-Jun expression and increases peripheral nerve myelination suggesting that inhibition of myelination is lifted. Restoring the Schwann cell phenotype will enable more WT P0 from the normal allele to reach myelin and contribute to clinical improvement. Supporting this explanation is the fact that we have recently demonstrated that patients who are haploinsufficient of *MPZ* have only a mild, late onset neuropathy, much milder than the neuropathy caused by R98C or Ser63del *MPZ* mutations [28].

### C3-PMP22 mice show altered proteostasis

The overexpression of PMP22 in animal models is sufficient to cause neuropathy, suggesting that increased dosage of PMP22 is the main contributing factor in CMT1A pathology [29, 58]. How the extra copy of *PMP22* causes disease is not fully understood, but studies in rodent models as well as in CMT1A patient derived fibroblast show that PMP22 can form cytosolic aggregates accompanied by reduced proteasomal activity, which may ultimately lead to cytotoxicity [17, 35, 45], PMP22 is a highly metastable protein, with more than

80% of newly synthesized PMP22 rapidly degraded by the proteasome, via ER-associated degradation (ERAD) [48, 49]. Recent *in vitro* work suggested that in normal condition, the levels of expression of PMP22 are close to the saturation capacity of the ERQC systems, and that overexpression of PMP22 leads to a disproportionate increase in misfolding and mis-trafficking [40]. Accordingly, examination of sciatic nerve from C22 mice, which show a severe dysmyelinating neuropathy, revealed increased expression of UPR markers such as BiP and CHOP [23]. In human, levels of PMP22 protein in CMT1A patient skin biopsies are elevated, even though the levels are more variable than in non-CMT1A samples [31, 37]. High levels of PMP22 protein have also been reported in sural nerve biopsies from CMT1A patients [19, 64]. Although the exact mechanism underlying the pathogenicity associated with expression of third copy of *PMP22* is not yet clearly defined, it is well established that a correct stoichiometry of PMP22 protein is required to maintain compact myelin integrity [27, 30]. Correction of PMP22 expression level reverses demyelination phenotype in a transgenic animal model [54] and the relevance of reducing PMP22 expression has been acknowledged as one potential therapeutic approach for CMT1A [47]. Recently, several approaches aimed at reducing PMP22 mRNA expression, such as anti-sense oligonucleotides (ASO), siRNA and AAV2/9-mediated shRNA targeting PMP22 have been successfully tested in murine models [6, 7, 20, 73], but their translation to humans still poses several hurdles.

We reasoned that a pharmacologic approach like IFB-088 aimed at attenuating general protein translation, thus reducing also PMP22 levels, would currently represent a more viable option than gene therapies. For this study we used the C3-PMP22 mice, that carry 3-4 copies of the human *PMP22* gene, and that is thought to represent the more appropriate model to study CMT1A, as compared to C22 mice (carrying 7-8 copies) [65]. We showed that in C3-PMP22 mice PMP22 overexpression results in the activation of the stress sensor BiP and of the PERK/P-eIF2_α_ pathway of the UPR, which supports the idea that activation of a stress response could be a contributing factor in CMT1A pathogenesis.

### IFB-088 treatment improves disease features in CMT1A mice

Treatment with IFB-088 improved motor capacity, neurophysiology, peripheral nerve morphology, and partially readjusted both myelin protein stoichiometry and stress levels in C3-PMP22 CMT1A mice. Importantly the treatment was started at PND15, when morphological deficits are already evident and ER-stress already activated, indicating that the treatment is ameliorating the disease and not simply preventing it. Intriguingly, some of these improvements were more prominent (treadmill) or present exclusively (neurophysiology) in females, whereas the improvement in dysmyelination was detectable in both sex although, again, some morphological parameters were ameliorated more in females than in males. The reasons for these differences are not clear. We ruled out a difference in IFB-088 exposure in males and females, as, in C57BL/6J mice, the same background as the C3-PMP22, no difference in term of IFB-088 pharmacokinetic profile was observed (Additional File 1: Supplementary Fig. 7). Moreover, we did not detect significant gender differences in the response to treatment in *Mpz*R98C/+ mice (this study), nor in *Mpz*S63del mice [12]. These results suggest that the difference may be more specific to the C3-PMP22 model itself. Of note, differences in motor capacity between genders have already been observed in different experimental setting in C3-PMP22 mice (F. Baas, *unpublished*) and in rotarod in the C22 model, with males performing significantly worse than females [51].

Neurophysiology analysis revealed an amelioration in nerve conduction velocity exclusively in females. The strong neurophysiological impairment in C3-PMP22 mice was accompanied by a complex and severe dysmyelinating phenotype, characterized by hypermyelination of small fibers and almost complete amyelination of large fibers, consistent with previous reports [65]. While myelination of large fibers improved almost equally in males and females after treatment with IFB-088, small fibers hypermyelination was more efficiently corrected in females. Whether this difference in morphological rescue is sufficient to explain the discrepancy in NCV after treatment remains unclear however as the speed of NCV depends predominantly on large diameter fibers [34].

The dysmyelinating phenotype in C3-PMP22 mice was accompanied by an increase in the levels of the ER-stress marker BiP and of P-eIF2_α_. Conversely, *Xbp1s* levels did not change significantly between WT and C3-PMP22 nerves, indicating that, unlike the CMT1B models *Mpz*R98C/+ and *Mpz*S63del, not all the UPR pathways were activated by PMP22 overexpression. This difference is likely related to different mechanisms of stress activation in the two models: whereas R98C and Ser63del P0 are misfolded proteins retained in the ER [53, 56, 70] where they activate a canonical UPR, overexpression of PMP22 is thought to overwhelm the ERAD-proteasome system, and the activation of ER-stress is likely a secondary event. In this respect it has been shown that ERAD impairment in Schwann cells results in ER-stress activation [68, 72]. We hypothesize that by prolonging the phosphorylation of eIF2_α_, IFB-088 attenuates the translational of highly expressed myelin proteins including PMP22. As a result, stress levels were restored towards WT levels in treated C3-PMP22 mice, and there was a partial readjustment of protein stoichiometry in nerves from both male and female mice.

Despite these very promising results, the C3-PMP22 was only partially improved, and not fully rescued, probably because, as for *MPZ*R98C/+ mice, the dysmyelinating phenotype of these mice is very severe to start with. Several questions remain to be answered, including whether a prolonged treatment beyond 4-month of age would further improve the phenotype, and whether a treatment initiated in adult mice would still provide benefits. Moreover, it should be noted that while the C3-PMP22 mouse represents a good model of CMT1A, currently there is no authentic animal model for this disease, as all available models express extra copies of the *PMP22* gene but without replicating the 1.4Mb duplication found in humans.

In summary, we have demonstrated that IFB-088 treatment improved the neuropathy in *Mpz*R98C/+ mice, a CMT1B mouse model, and in C3-PMP22 mice, a CMT1A mouse model. This is the second example of IFB-088’s ability to improve the neuropathy in a mouse model of CMT1B. We have identified many additional mutations in *MPZ* that activate the UPR [4] and expect that IFB-088 may prove beneficial in many of these patients. In addition, our study confirms a role of ER-stress in the pathogenesis of CMT1A and demonstrate the ability of IFB-088 to assist the ER-stress response to the over-expression of PMP22 and potentially treat patients with CMT1A. Thus, IFB-088, which completed successfully phase I clinical trial, represents a new and promising therapeutic option for CMT1A and CMT1B. Due to his mode of action, IFB-088 has the unique potential to provide benefits to different CMT subtypes caused by different gene defects. Moreover, the benefits of IFB-088 have been also demonstrated in Amyotrophic Lateral Sclerosis and in Multiple Sclerosis models [8, 9, 12], suggesting that managing UPR and ER-stress are promising strategies for multiple neurodegenerative diseases.

## Abbreviations

AD: autosomal dominant
AR: autosomal recessive
ASO: anti-sense oligonucleotides
ATF4: activating transcription factor 4
BiP: immunoglobulin heavy chain-binding protein
CHOP: C/EBP homologous protein
CMT: Charcot Marie Tooth
CMAP: compound muscle action potential
DRG: dorsal root ganglia
eIF2α: alpha subunit of the eukaryotic translation initiation factor 2
ER: endoplasmic reticulum
ERAD: ER-associated degradation
ERQC: endoplasmic reticulum protein quality control
Grp94: glucose regulated protein 94
IRE1: inositol requiring enzyme 1
MBP: myelin binding protein
MPZ: myelin protein zero
MNCV: motor nerve conduction velocity
NCV: nerve conduction velocity
NF: neurofilament
PERK: protein-kinase RNA-like endoplasmic reticulum kinase
PMP22: peripheral myelin protein 22
PND: post-natal day
PPP1R15A: Protein Phosphatase 1 Regulatory Subunit 15A
PQC: protein quality control
SNAP: sensory nerve action potential
SNCV: sensory nerve conduction velocity
UPR: unfolded protein response
Tr^J^: Trembler J
TEM: transmission electron microscopy
WB: western blot
WT: wild type
XBP1: X-box-binding protein-1.

## Declarations

### Availability of data and materials

The data that support the findings of this study are available from the corresponding author, upon reasonable request.

### Authors’ contributions

M.D., M.E.S., P.G., P.M., C.T. contributed to the study conception and design. F.B. provided the C3-PMP22 mouse model. R.M. performed the myelinating DRG explant cultures. Y.B. and DW performed the study in the MpzR98C/+ mouse model. V.G.V., C.S., C.F., R.M., T.T., F.F., performed the study in the C3-PMP22 mouse model. F.B., U.D. performed the neurophysiological evaluation of the C3-PMP22 mice. M.D., M.E.S. and C.T. wrote the manuscript. All authors commented on previous versions of the manuscript. All authors read and approved the final manuscript.

## Acknowledgements

We thank the Charcot-Marie-Tooth Association for their continuous support. We are grateful to Dr. Patrizia D’Adamo and the Mouse Behavior Facility at IRCCS Ospedale San Raffaele. Part of this work was carried out in ALEMBIC, an advanced microscopy laboratory established by IRCCS Ospedale San Raffaele and Università Vita-Salute San Raffaele. M.D. is supported by Fondazione Telethon (GGP19099), CMTA (Charcot-Marie-Tooth Association, USA) and CMTRF (Charcot-Marie-Tooth Research Foundation).

---

**Additional file 1:** Supplementary figures S1 - S7.

