## Supplementary Figures for "Treatment with IFB-088 improves neuropathy in CMT1A and CMT1B mice"

### Additional file 1

### Supplementary figures legends

**Supplementary Figure 1: IFB-088 improves myelination in DRG explants from *Mpz*S63del mice.** Dorsal root ganglia (DRG) were dissected from embryos (E13.5) of *Mpz*S63del mice and *Mpz*R98C/+ mice. The myelination process was induced with ascorbic acid. After 2-week of treatment with vehicle or the indicated concentration of IFB-088, the DRGs were fixed and nuclei visualized by DAPI staining; axons and myelin were visualized by immunostaining with neurofilament antibody (NF) and with myelin basic protein (MBP) antibodies respectively. (**a**) Representative pictures and number of myelinated internodes per field, expressed in % of WT, in WT and *Mpz*R98C/+ DRG explant cultures treated with vehicle for 2 weeks. (**b**) Representative pictures and number of myelinated internodes per field, expressed in % of WT, in WT and *Mpz*S63del DRG explant cultures treated with vehicle for 2 weeks. (**c**) Representative pictures. (**d**) Number of myelinated internodes per field in *Mpz*S63del DRG explant cultures treated with vehicle or the indicated concentration of IFB-088 for 2 weeks. Mean ±SEM. $n$=3 independent experiments. *$P$<0.05 One-way ANOVA followed by Friedman's test.

**Supplementary Figure 2: IFB-088 treatment does not impact WT and *Mpz*R98C/+ mice body weight.** Body weight of WT and *Mpz*R98C/+ mice treated with vehicle *b.i.d.* or IFB-088 1 mg/kg *b.i.d.* expressed in % of body weight over 20-week treatment periods ($n$=19-36 mice per condition).

**Supplementary Figure 3: Impact of 3-month IFB-088 treatment on motor function and nerve conduction velocity in *Mpz*R98C/+ mice.** (**a**) Diagram of the treatment strategy. 30-day-old WT and *Mpz*R98C/+ mice were orally administered with vehicle *b.i.d.* or IFB-088 1mg/kg *b.i.d.* for 3 months. (**b**) Four limb grip strength max values average of 10 trials. Data were expressed in grams (g) as mean ±SEM. *n*=11-19 mice per condition. (**c**) Rotarod analysis. Data are expressed in seconds (s) as mean ±SEM. *n*=10-19 mice per condition. (**d**) Motor nerve conduction velocity (MNCV). Data are expressed in meter/second (m/s) as mean ±SEM. *n*=10-24 mice per condition. (**e**) Sensory nerve conduction velocity (SNCV). Data are expressed in meter/second (m/s) as mean ±SEM. *n*=5-20 mice per condition. ***$P$<0.001, ****$P$<0.0001 by Student's T-test; #$P$<0.05, ##$P$<0.01 Mann-Whitney.

**Supplementary Figure 4: At PND15, C3-PMP22 mice present a CMT1A-like phenotype with a relative overexpression of PMP22 and expression of ER Stress/UPR markers.** (**a**) Toluidine-blue stained transverse semithin-sections from sciatic nerve from 15-day-old WT and C3-PMP22 (C3) mice. Scale bar, 10µm. (**b**) Left: Representative WB for PMP22 and P0 on sciatic nerve protein extracts from 15-day-old WT and C3-PMP22 (C3) mice. Tubulin and Coomassie staining of total proteins were used as loading controls. Right: Quantification of the PMP22/P0 protein ratio relative to total protein. #$P$<0.05 by Mann-Whitney. (**c**) Top: Representative WB for BiP, P-eIF2α and tubulin on sciatic nerve protein extracts from 15-day old WT and C3-PMP22 (C3) mice. Bottom: quantification of protein level from *n*=6 independent nerves. Tubulin was used as loading control. Student's T-test.

**Supplementary Figure 5: Treatment with IFB-088 improves C3-PMP22 mice sciatic nerve morphology.** Toluidine blue-stained sciatic nerve sections from (**a**) female and (**b**) male WT mice treated with vehicle *b.i.d.* and C3-PMP22 mice treated with vehicle or IFB-088 at 0.5 or 1mg/kg *b.i.d.* for 12 weeks. Scale bar, 10µm.

**Supplementary Figure 6: Treatment with IFB-088 improves morphology of C3-PMP22 quadriceps femoral nerve.** Scatter plot of quadriceps femoral nerve g-ratios from WT mice treated with vehicle *b.i.d.* and C3-PMP22 (C3) mice treated with vehicle *b.i.d.* or IFB-088 at 0.5 or 1mg/kg *b.i.d.* (**a**) Females, (**b**) males. Note the "cloud" of axons with diameter lower than 1µm mostly restricted to vehicle treated C3-PMP22 nerves, and the appearance of a considerable number of myelinated axons larger than 5-6µm in treated C3-PMP22 nerves. Percentage of myelinated axons per axons size. (**c**) Females, (**d**) males. *n*=2-4 nerves per condition.

**Supplementary Figure 7: IFB-088 pharmacokinetic profile.** Plasma IFB-088 concentration in C57BL6/J males and females after a single administration at 4 mg/kg. Data are expressed in ng/mL as a mean.

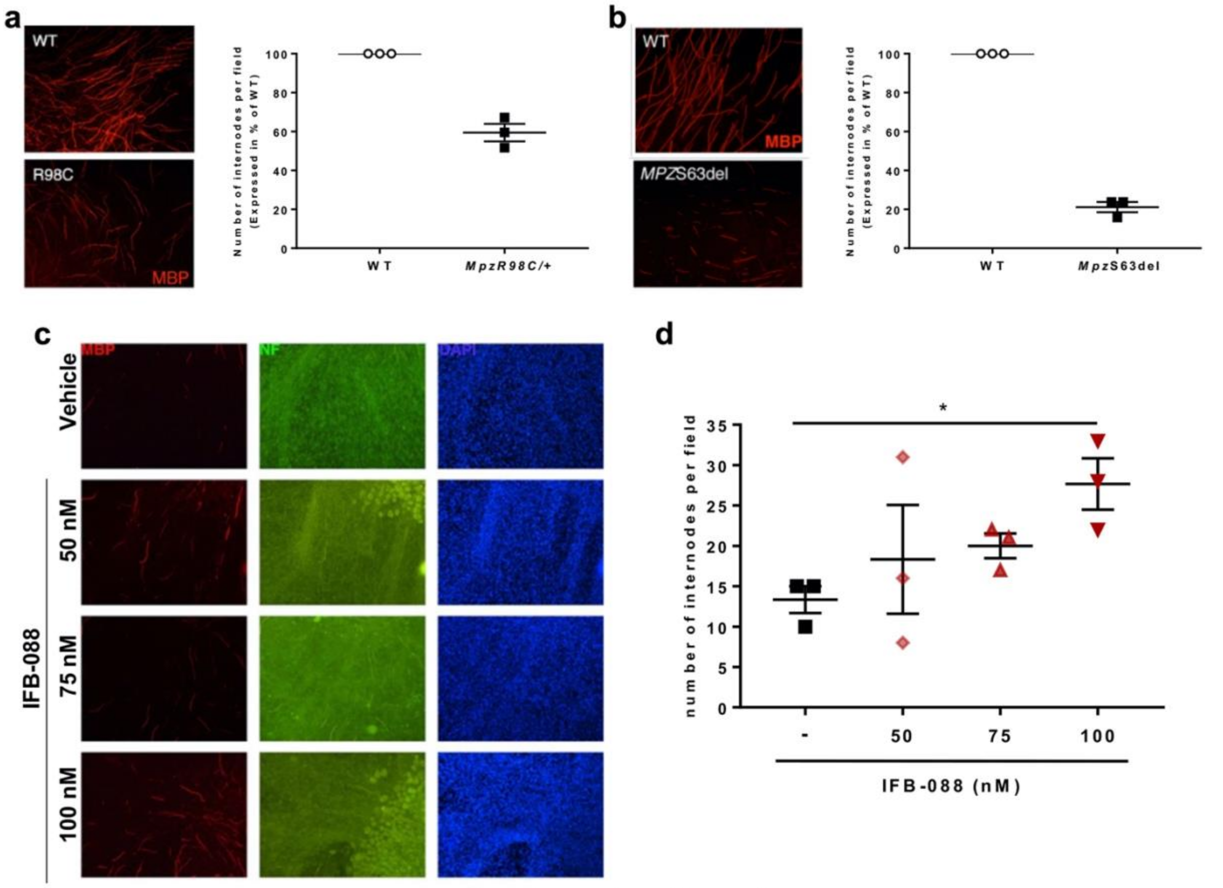


Supplementary Figure 1

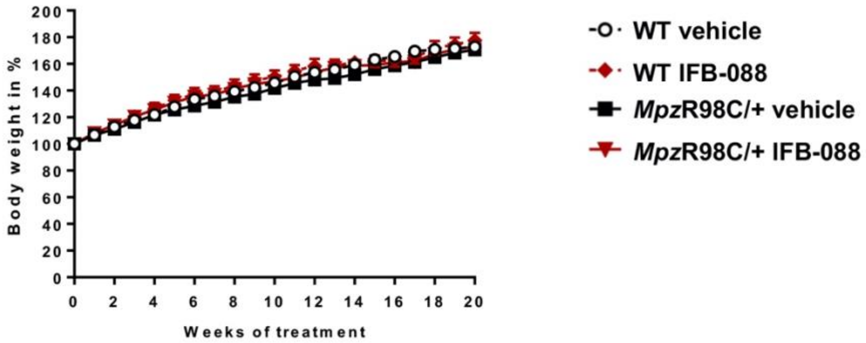


**Supplementary Figure 2**

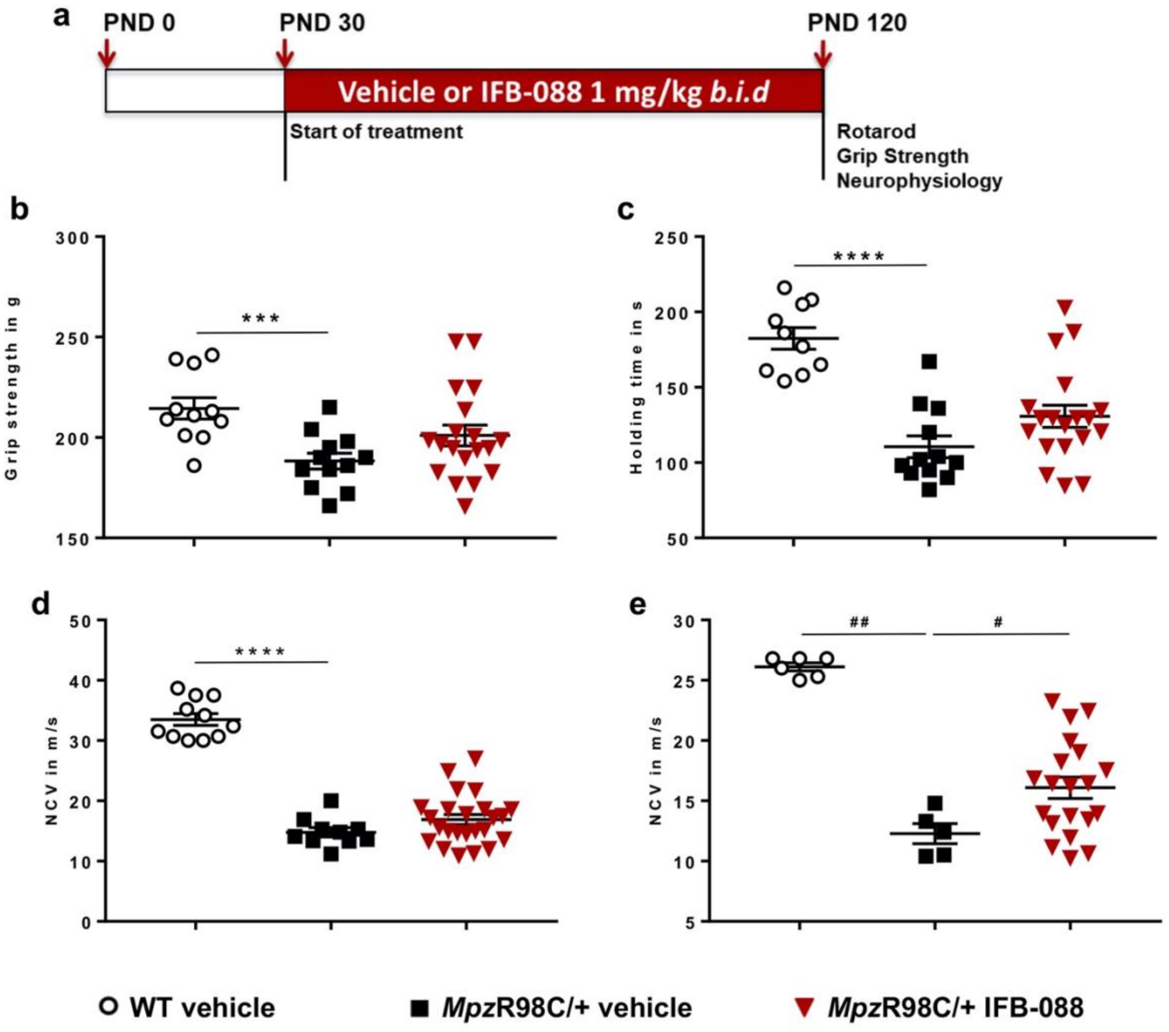


**Supplementary Figure 3**

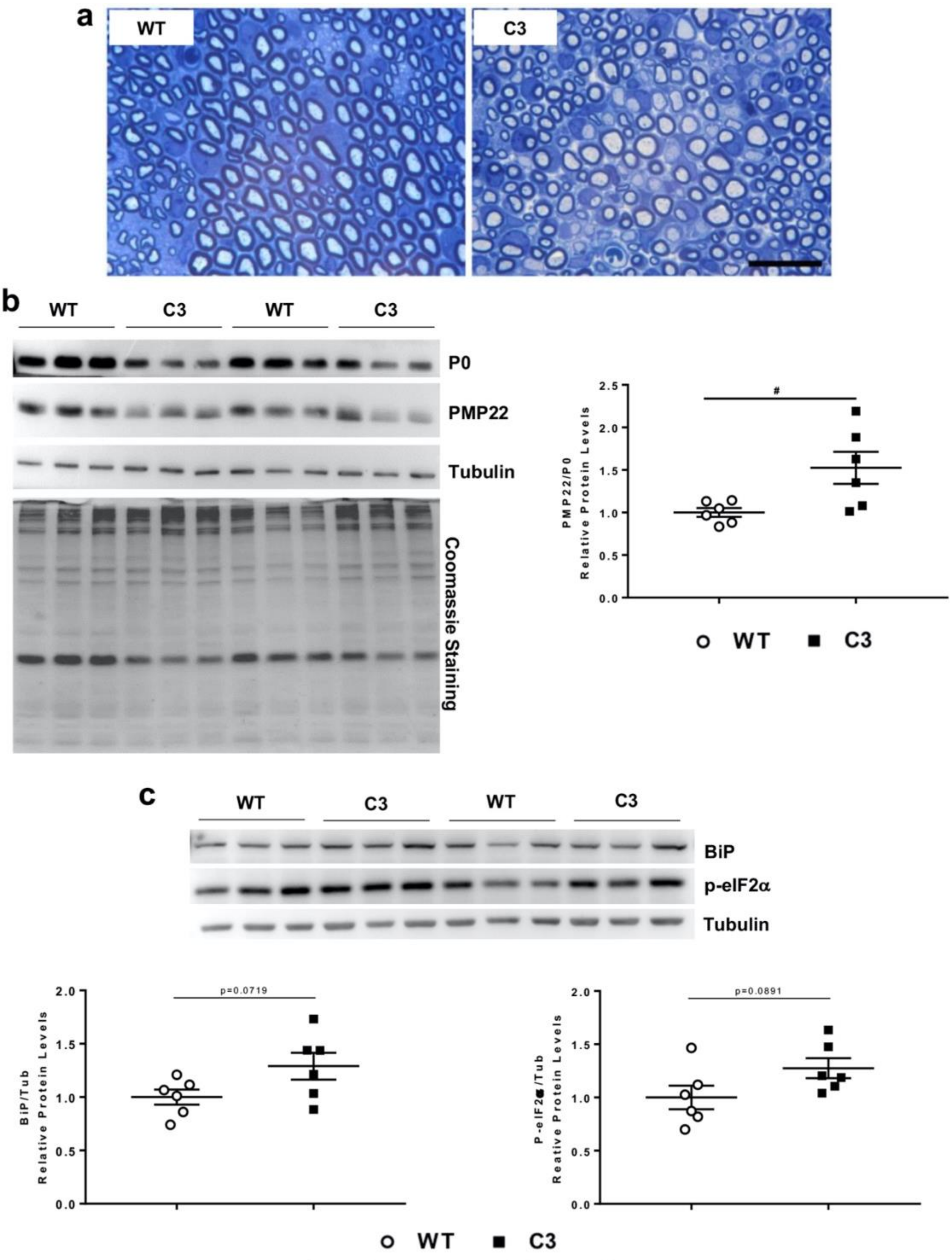


**Supplementary Figure 4**

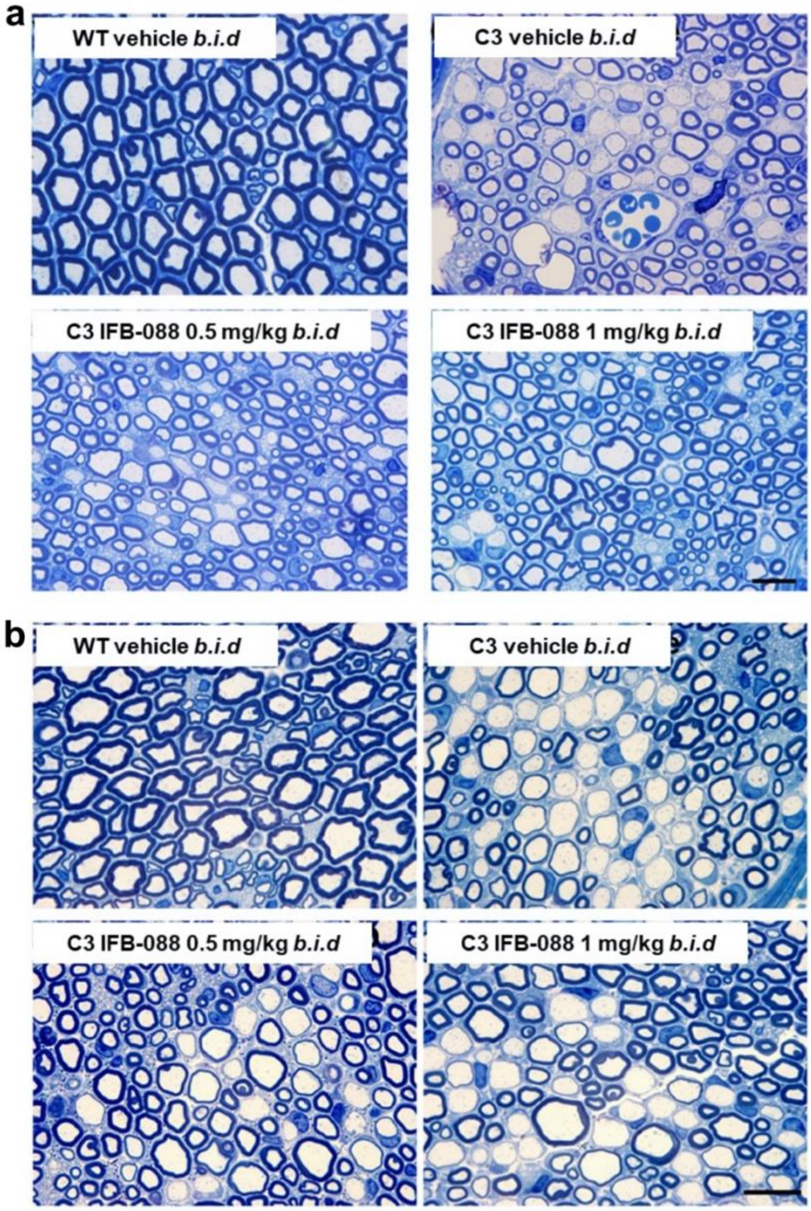


**Supplementary Figure 5**

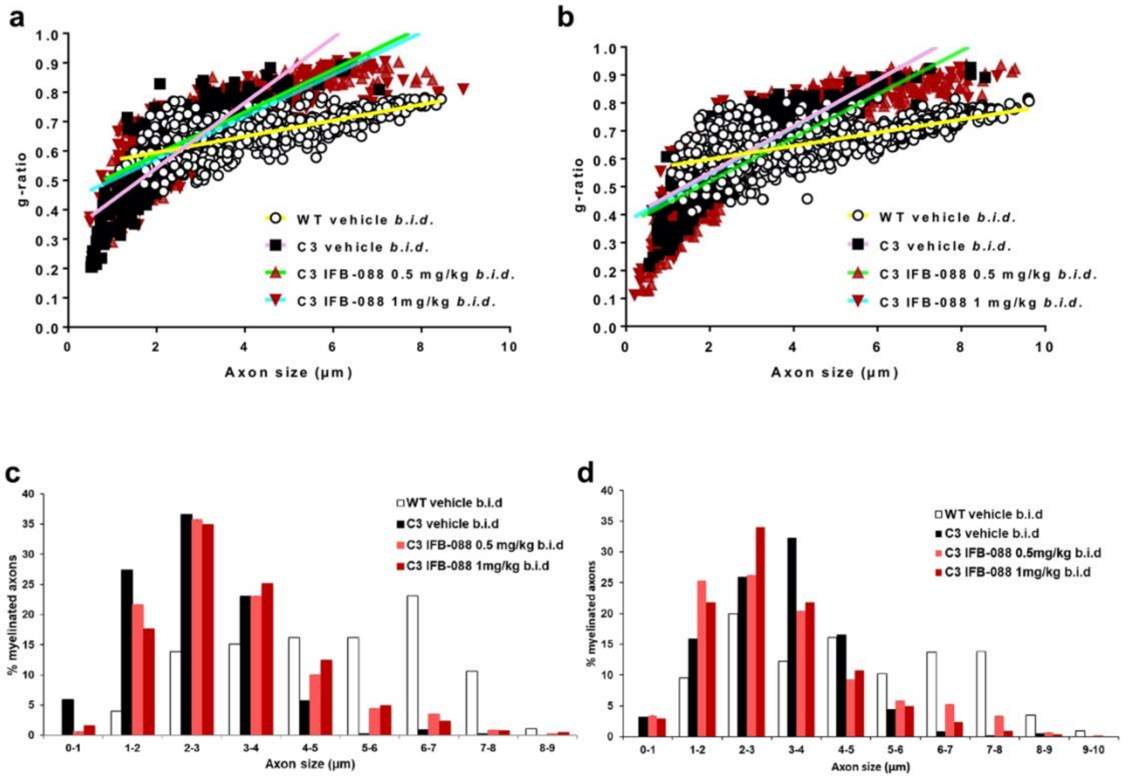


**Supplementary Figure 6**

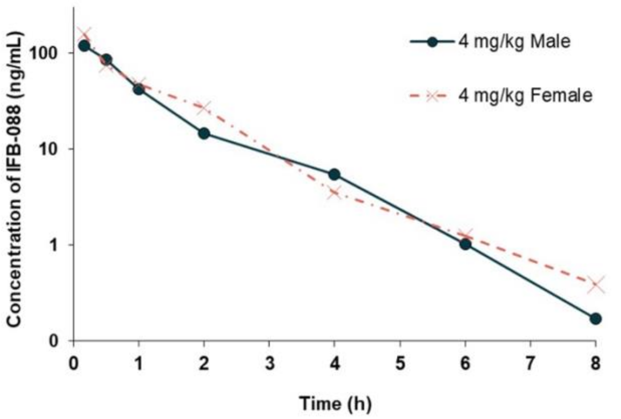


**Supplementary Figure 7**
